# A GPU-based computational framework that bridges Neuron simulation and Artificial Intelligence

**DOI:** 10.1101/2022.06.12.495784

**Authors:** Yichen Zhang, Gan He, Xiaofei Liu, J.J. Johannes Hjorth, Alexander Kozlov, Yutao He, Shenjian Zhang, Lei Ma, Jeanette Hellgren Kotaleski, Yonghong Tian, Sten Grillner, Kai Du, Tiejun Huang

## Abstract

Biophysically detailed multi-compartment models are powerful tools to explore computational principles of the brain and also serve as a theoretical framework to generate algorithms for artificial intelligence (AI) systems. However, the expensive computational cost severely limits the applications in both the neuroscience and AI fields. The major bottleneck during simulating detailed compartment models is the ability of a simulator to solve large systems of linear equations. Here, we present a novel **D**endritic **H**ierarchical **S**cheduling (DHS) method to markedly accelerate such process. We theoretically prove that the DHS implementation is computationally optimal and accurate. This GPU-based method performs at 2-3 orders of magnitude higher speed than that of the classic serial Hines method in the conventional CPU platform. We build a DeepDendrite framework, which integrates the DHS method and the GPU computing engine of the NEURON simulator and demonstrate applications of DeepDendrite in neuroscience and AI tasks. We investigated how spatial patterns of spine inputs affect neuronal excitability in a detailed human pyramidal neuron model with 25,000 spines; and examined how dendrites protect morphologically detailed neural networks against adversarial attacks in typical image classification tasks.

## Introduction

Deciphering the coding and computational principles of neurons is essential to neuroscience. Mammalian brains are composed of more than thousands of different types of neurons with unique morphological and biophysical properties. Even though it is no longer conceptually true, the “point-neuron” doctrine^1^, in which neurons were regarded as simple summing units, is still widely applied in neural computation, especially in neural network analysis. In recent years, modern artificial intelligence (AI) has utilized this principle and developed powerful tools, such as the artificial neural network (ANN)^2^. However, in addition to comprehensive computations at the single neuron level, subcellular compartments, such as neuronal dendrites, can also carry out nonlinear operations as independent computational units^3-7^. Furthermore, dendritic spines, small protrusions that densely cover dendrites in spiny neurons, can compartmentalize synaptic signals, allowing them to be separated from their parent dendrites *ex vivo* and *in vivo*^8-11^.

Simulations using biologically detailed neurons provide a theoretical framework for linking biological details to computational principles. The core of the biophysically detailed multi-compartment model framework^12, 13^ allows us to model neurons with realistic dendritic morphologies, intrinsic ionic conductance, and extrinsic synaptic inputs. The backbone of the detailed multi-compartment model, i.e., dendrites, is built upon the classical Cable theory^12^, which models the biophysical membrane properties of dendrites as passive cables, providing a mathematical description of how electronic signals invade and propagate throughout complex neuronal processes. By incorporating Cable theory with active biophysical mechanisms such as ion channels, excitation, inhibition, etc., a detailed multi-compartment model can achieve cellular and subcellular neuronal computations beyond experimental limitations^4, 7^.

In addition to its profound impact on neuroscience, biologically detailed neuron models recently were utilized to bridge between neuronal structural and biophysical details and AI. The prevailing technique in the modern AI field is ANNs consisting of point-neurons, an analog to biological neural network. Although ANNs with “backpropagation-of-error” (backprop) algorithm achieve remarkable performance in specialized applications, even beating top human professional players in the games of Go and chess^14, 15^, the human brain still outperforms ANNs in other domains, for example, general intelligence. Recent theoretical studies suggest that dendritic integration is crucial in generating efficient learning algorithms that potentially exceed backprop in parallel information processing^16-18^. Furthermore, a single detailed multi-compartment model can learn network-level nonlinear computations for point-neurons by adjusting only the synaptic strength^19, 20^, demonstrating the full potential of the detailed models in building more powerful brain-like AI systems. Therefore, it is of high priority to expand paradigms in brain-like AI from single detailed neuron models to large-scale biologically detailed networks.

One long-standing challenge of the detailed simulation approach lies in its exceedingly high computational cost, which has severely limited its application to neuroscience and AI. The major bottleneck of the simulation is to solve linear equations based on the foundational theories of detailed modeling^12, 21, 22^. To improve the efficiency, the classic Hines method reduces the time complexity for solving equations from O(n^3^) to O(n), which has been widely applied as the core algorithm in popular simulators such as NEURON^23^ and GENESIS^24^. However, this method uses a serial approach to process each compartment sequentially. When a simulation involves multiple biophysically detailed dendrites with dendritic spines, the linear equation matrix (‘Hines Matrix’) scales accordingly with an increasing number of dendrites or spines (Fig. 1e), making Hines method no longer practical, since it poses a very heavy burden on the entire simulation.

**Figure 1.**
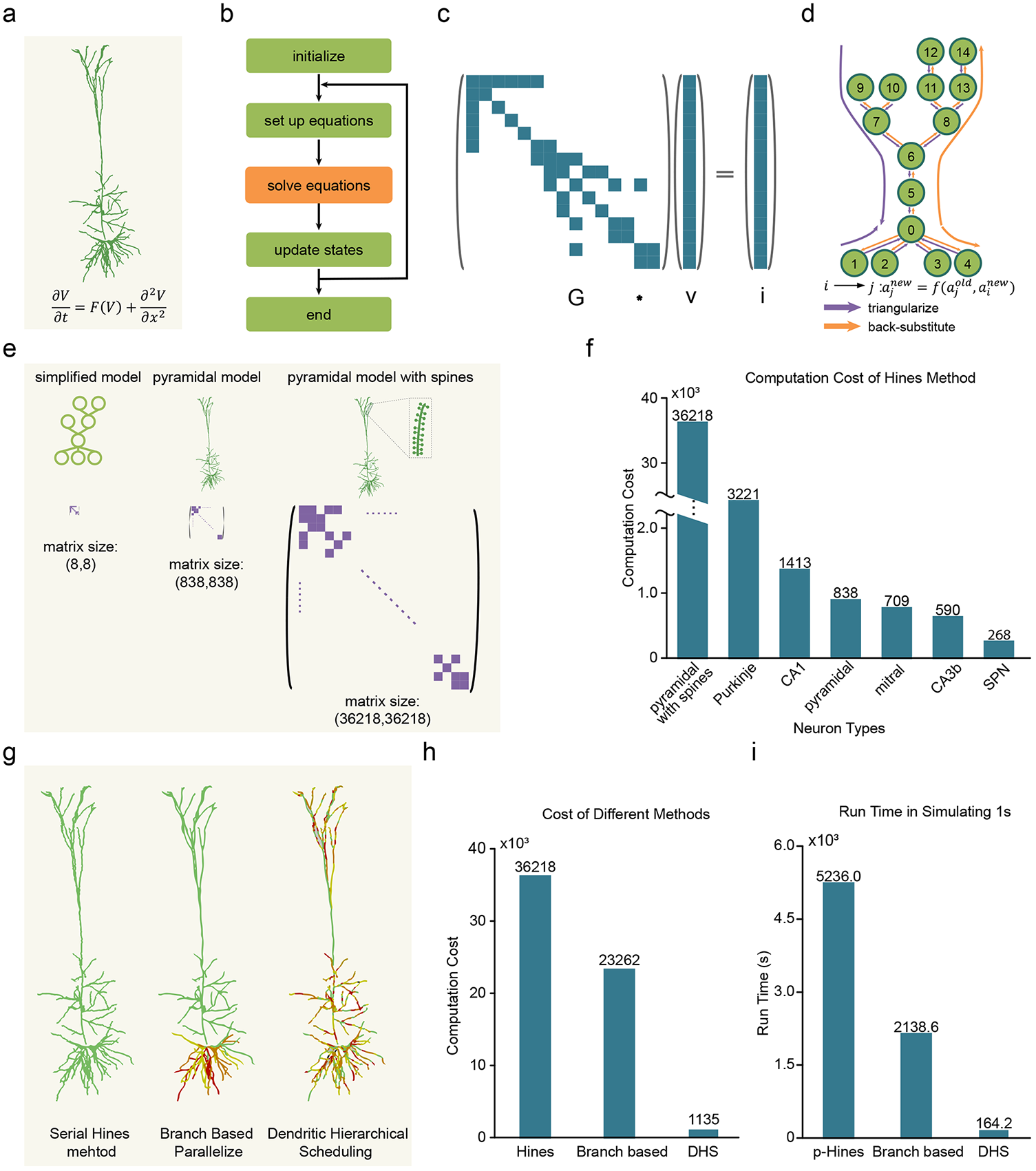
Accelerate simulation of biophysically detailed neuron models. **a**, A reconstructed layer-5 pyramidal neuron model and the mathematical formula used with detailed neuron models. **b**, Workflow when numerically simulating detailed neuron models. The equation solving phase is the bottleneck in the simulation. **c**, An example of linear equations in the simulation. **d**, Data dependency of the Hines method when solving linear equations in **c**. **e**, The size of Hines matrix scales with model complexity. The number of linear equations system to be solved undergoes a significant increase when models are growing more detailed. **f**, Computational cost of the serial Hines method on different types of neuron models. **g**, Illustration of different solving methods. Different parts of a neuron are assigned to multiple processing units in parallel methods (mid, right), shown with different colors. In serial method (left), all compartments are computed with one unit. **h**, Computational cost of three methods in **g** when solving equations of a pyramidal model with spines. **i**, Run time of different methods on solving equations for 500 pyramidal models with spines. The run time indicates the time consumption of 1s simulation (solving the equation 40000 times during the simulation). p-Hines: parallel method in CoreNEURON (on GPU); Branch based: branch based parallel method (on GPU); DHS: Dendritic hierarchical scheduling method (on GPU).

During past decades, tremendous progress has been achieved to speed up the Hines method by using parallel methods at a cellular level, which enables to parallelize the computation of different parts in each cell^25-30^. However, current cellular level parallel methods often lack an efficient parallelization strategy or lack sufficient numerical accuracy as compared to the original Hines method.

Here, we develop a fully automatic, numerically accurate, and optimized simulation tool that can significantly accelerate computation efficiency and reduce computational cost. In addition, this simulation tool can be seamlessly adopted for establishing and testing neural networks with biological details for machine learning and AI applications. Critically, we formulate the parallel computation of the Hines method as a mathematical scheduling problem and generate a Dendritic Hierarchical Scheduling (DHS) method based on combinatorial-optimization^31^ and parallel-computing theory^32^. We demonstrate that our algorithm provides an optimal scheduling without any loss of precision. Furthermore, we have optimized DHS for the currently most advanced GPU chip by leveraging the GPU memory hierarchy and memory accessing mechanisms. Together, DHS can speed up computation 60-1,500 times (Table S1) compared to the classic simulator NEURON^23^ while maintaining identical accuracy.

To enable detailed dendritic simulations for use in AI, we next establish the DeepDendrite framework by integrating the DHS-embedded CoreNEURON (an optimized compute engine for NEURON) platform^33^ as the simulation engine and two auxiliary modules (I/O module and learning module) supporting dendritic learning algorithms during simulations. DeepDendrite runs on the GPU hardware platform, supporting both regular simulation tasks in neuroscience and learning tasks in AI.

Last but not least, we also present several applications using DeepDendrite, targeting a few critical challenges in neuroscience and AI: (1) We demonstrate how spatial patterns of dendritic spine inputs affect neuronal activities with neurons containing spines throughout the dendritic trees (full-spine models). DeepDendrite enables us to explore neuronal computation in a simulated human pyramidal neuron model with ∼25,000 dendritic spines. (2) We establish an artificial neural network with morphologically detailed human pyramidal neurons and train this network to perform typical image classification tasks. DeepDendrite reduces the training time from days to minutes, which enables detailed network models to perform AI tasks. We further explore previously proposed hypothesis about the roles of the dendrites in defending against adversarial attacks^34^, an intentionally designed perturbation to deceive ANNs but nearly unperceivable to human vision. Models with morphologically detailed dendrites show superior performance with strong robustness in defending against adversarial attacks.

All source code for DeepDendrite, the full-spine models and the detailed dendritic network model are publicly available online (see Method). Our open-source learning framework can be readily integrated with other dendritic learning rules, such as learning rules for nonlinear (full-active) dendrites^19^, burst-dependent synaptic plasticity ^18^, and learning with spike prediction^35^. Overall, our study provides a complete set of tools that have the potential to change the current computational neuroscience community ecosystem. By leveraging the power of GPU computing, we envision that these tools will facilitate system-level explorations of computational principles of the brain’s fine structures, as well as promote the interaction between neuroscience and modern AI.

## Results

Computing ionic currents and solving linear equations are two critical phases when simulating biophysically detailed neurons, and which are time-consuming and poise severe computational burden. Fortunately, computing ionic currents of each compartment is a fully independent process so that it can be naturally parallelized on devices with massive parallel-computing units like GPUs^36^. As a consequence, solving linear equations becomes the remaining bottleneck for the parallelization process (Fig. 1a-f).

To tackle this bottleneck, cellular level parallel methods have been developed, which accelerate single-cell computation by “splitting” a single cell into several compartments that can be computed in parallel^25, 26, 37^. However, such methods rely heavily on prior knowledge in order to generate practical strategies on how to split a single neuron into compartments (Fig. 1g-i, S1). Hence, it becomes less efficient for neurons with asymmetrical morphologies, e.g., pyramidal neurons and Purkinje neurons.

### Dendritic Hierarchical Scheduling (DHS) Method

Our aim is to develop a more efficient and precise parallel method for simulation of biologically detailed neural networks. First, we establish the criteria for the accuracy of a cellular level parallel method. Based on the theories in parallel computing^32^, we propose three conditions to make sure a parallel method will yield identical solutions as the serial computing Hines method according to the data dependency in Hines method (see Methods). Then to theoretically evaluate the run time, i.e. efficiency, of the serial and parallel computing methods, we introduce and formulate the concept of computational cost as the number of steps a method takes in solving equations (see Methods).

Based on the simulation accuracy and computational cost, we formulate the parallelization problem as a mathematical scheduling problem (details are in Methods). In simple terms, we view a single neuron as a tree with many nodes (compartments). For k parallel threads, we can compute at most k nodes at each step, but we need to ensure a node is computed *only if* all its children nodes have been processed; our goal is to find a strategy with the minimum number of steps for the entire procedure.

To generate an optimal partition, we propose a method called Dendritic Hierarchical Scheduling (DHS) (theoretical proof is presented in the Methods). The key idea of DHS is to prioritize deep nodes (Fig 2a), which results in a hierarchical schedule order. The DHS method includes two steps: analyzing dendritic topology and finding the best partition: (1) Given a detailed model, we first obtain its corresponding dependency tree and calculate the depth of each node (the depth of a node is the number of its ancestor nodes) on the tree (Fig 2 b,c). (2) After topology analysis, we search the candidates and pick at most *k* deepest candidate nodes (a node is a candidate only if all its children nodes have been processed). This procedure repeats until all nodes are processed (Fig. 2d).

**Figure 2.**
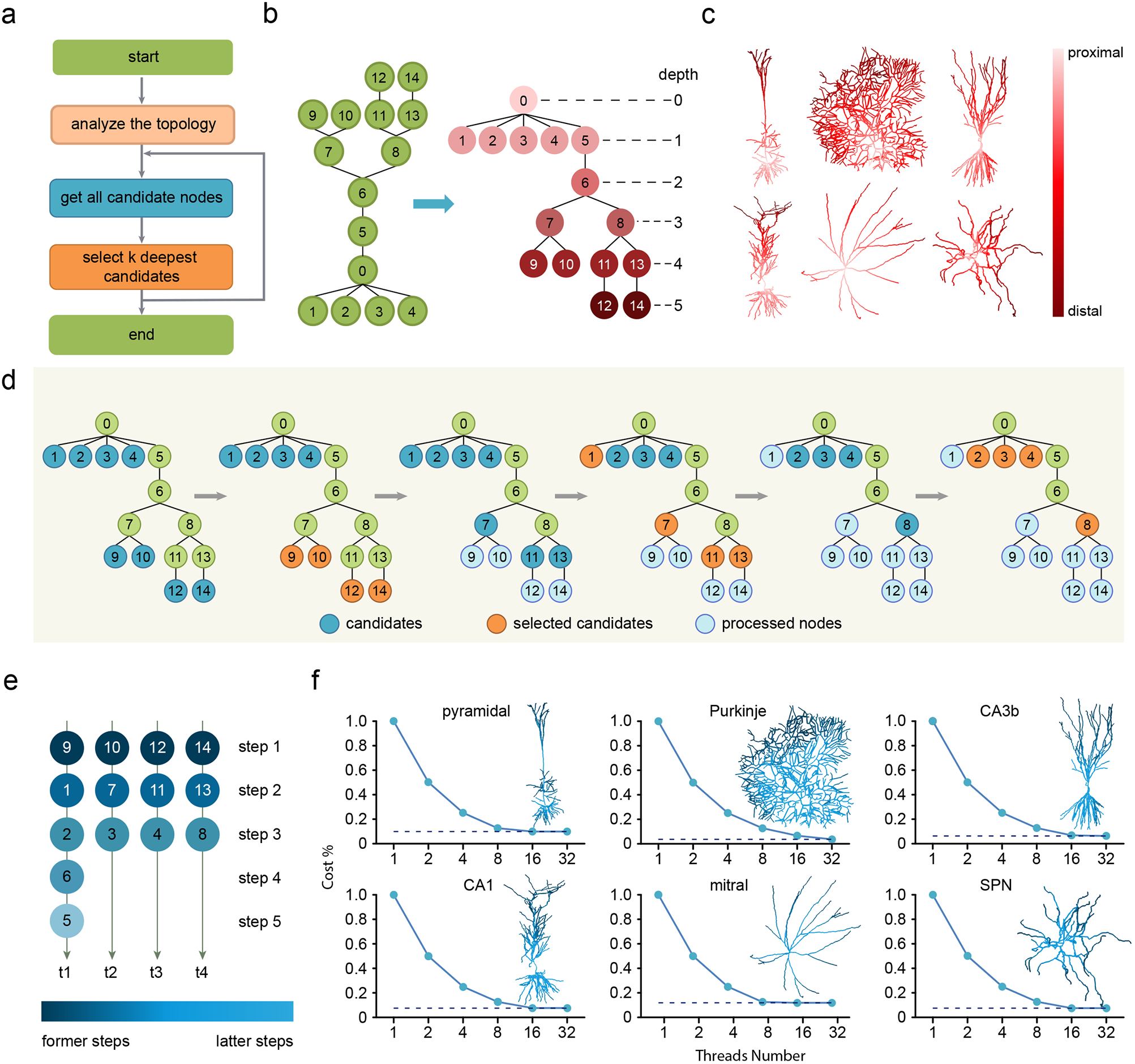
Dendritic Hierarchical Scheduling (DHS) method significantly reduce the computational cost, i.e. computational steps in solving equations. **a**, DHS work flow. DHS processes k deepest candidate nodes each iteration. **b**, Illustration of calculating node depth of a compartmental model. The model is first converted to a tree structure then the depth of each node is computed. Colors indicate different depth values. **c**, Topology analysis on different neuron models. Six neurons with distinct morphologies are shown here. For each model, soma is selected as root of the tree so the depth of the node increases from the soma (0) to the distal dendrites. **d**, Illustration of performing DHS on the model in **b** with four threads. Candidates: nodes that can be processed. Selected candidates: nodes that are picked by DHS, i.e. the k deepest candidates. Processed nodes: nodes that have been processed before. **e**, Parallelization strategy obtained by DHS after the process in **d**. Each node is assigned to one of the four parallel threads. DHS reduces the steps of serial node processing from 14 to 5 by distributing nodes to multiple threads. **f**, Computational cost (steps taken in the equation solving phase) when applying DHS with different thread numbers on different types of models. Relative cost, i.e. the ratio of DHS cost to NEURON’s serial Hines method cost, is used here.

Take a simplified model with 15 compartments as an example, using the serial computing Hines method, it takes 14 steps to process all nodes, while using DHS with four parallel units can partition its nodes into five subsets (Fig. 2d): {{9,10,12,14}, {1,7,11,13}, {2,3,4,8}, {6}, {5}}. Because nodes in the same subset can be processed in parallel, it takes only five steps to process all nodes using DHS (Fig. 2e).

Next, we apply the DHS method on six representative detailed neuron models (selected from ModelDB, https://senselab.med.yale.edu/ModelDB) with different numbers of threads (Fig. 2f):, including cortical and hippocampal pyramidal neurons^39-41^, cerebellar Purkinje neurons^42^, striatal spiny-projection neurons (SPN^43^), and olfactory bulb mitral cells^44^, covering the major principle neurons in sensory, cortical and subcortical areas. We then measured the computational cost. The relative computational cost here is defined by the proportion of the computational cost of DHS to that of the serial Hines method. The computational cost, i.e. the number of steps taken in solving equations, drops dramatically with increasing thread numbers. For example, with 16 threads, the computational cost of DHS is 7%-10% as compared to the serial Hines method. Intriguingly, the DHS method reaches the lower bounds of their computational cost for presented neurons when given 16 or even 8 parallel threads (Fig. 2f), suggesting adding more threads does not improve performance further because of the dependencies between compartments.

Together, we generate a DHS method that enables automated analysis of the dendritic topology and optimal partition for parallel computing. It is worth noting that DHS finds the optimal partition before simulation starts, and no extra computation is needed to solve equations.

### Speeding Up DHS by GPU Memory Boosting

DHS computes each neuron with multiple threads, which consumes a vast amount of threads when running neural network simulations. Graphics Processing Units (GPU) consists of massive processing units (streaming processor, SP, Fig. 3a,b) for parallel computing^45^. In theory, many SPs on the GPU should support efficient simulation for large-scale neural networks (Fig. 3c). However, we consistently observed that the efficiency of DHS significantly decreased when the network size grew, which might result from scattered data storage or extra memory access caused by loading and writing intermediate results (Fig. 3d, left).

**Figure 3.**
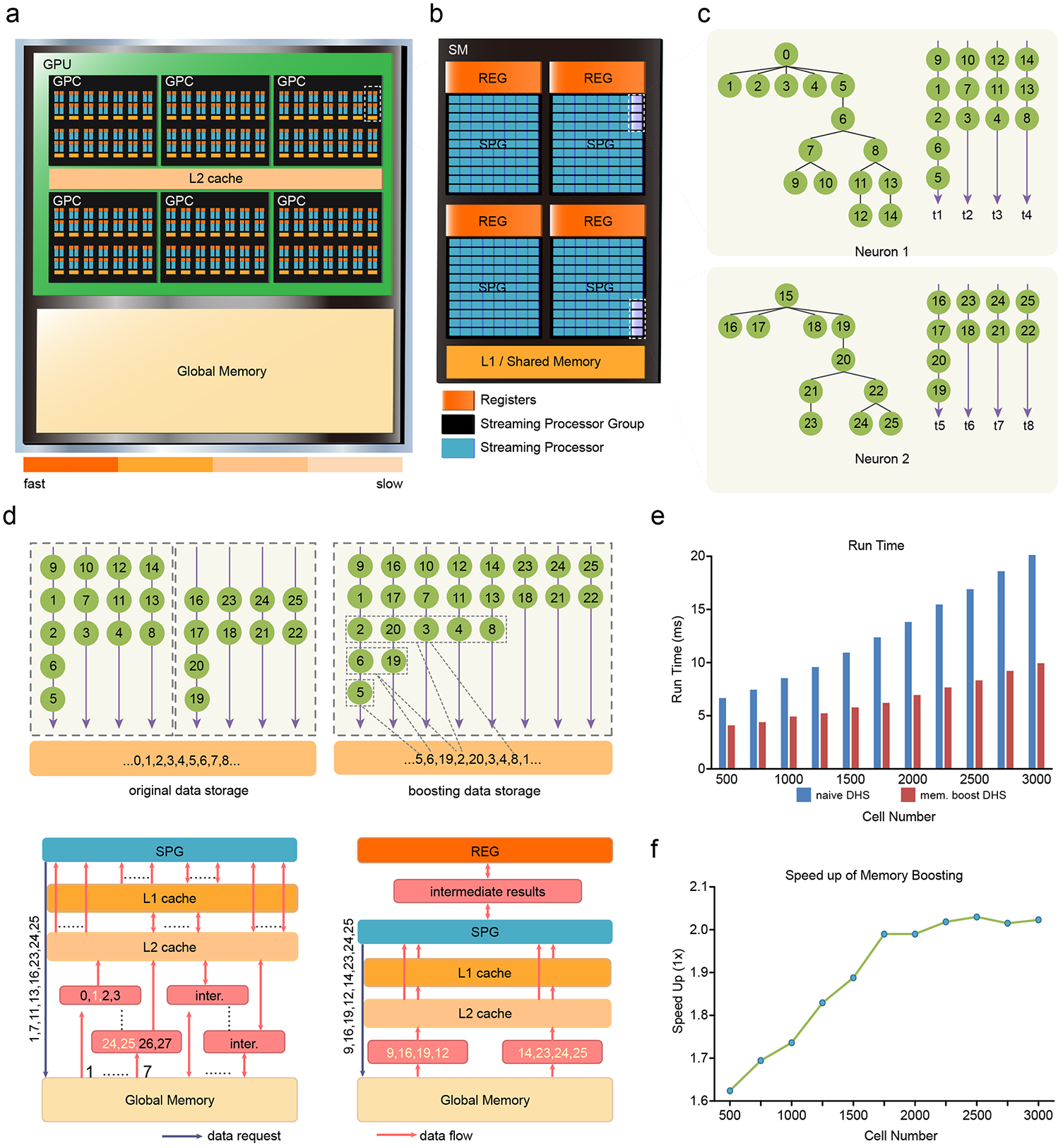
GPU memory boosting further accelerates DHS. **a**, GPU architecture and its memory hierarchy. Each GPU contains massive processing units (stream processors). Different types of memory have different throughput. **b**, Architecture of Streaming Multiprocessors (SMs). Each SM contains multiple streaming processors, registers and L1 cache. **c**, Apply DHS on two neurons, each with four threads. During simulation, each thread executes on one stream processor. **d**, Memory optimization strategy on GPU. Top pannel, thread assignment and data storage of DHS, before (left) and after (right) memory boosting. Bottom, an example of single step in triangularization when simulating two neurons in **d**. Processors send a data request to load data for each thread from global memory. Without memory boosting (left), it takes seven transactions to load all request data and some extra transactions for intermediate results. With memory boosting (right), it takes only two transactions to load all request data, registers are used for intermediate results, which further improve memory throughput. **e**, Run time of DHS (32 threads each cell) with and without memory boosting on multiple layer 5 pyramidal models with spines. **f**, Speed up of memory boosting on multiple layer 5 pyramidal models with spines. Memory boosting brings 1.6-2 folds speedup.

We solve this problem by GPU memory-boosting, a method to increase memory throughput by leveraging GPU’s memory hierarchy and access mechanism. Based on the memory loading mechanism of GPU, successive threads loading aligned and successively-stored data lead to a high memory throughput compared to accessing scatter-stored data, which reduces memory throughput^45, 46^. To achieve the high throughput, we first align computing orders of nodes and rearrange threads according to the number of nodes on them. Then we permute data storage in global memory, consistent with computing orders, i.e. nodes that are processed at the same step are stored successively in global memory. Moreover, we use GPU registers to store intermediate results, further strengthening memory throughput. The example shows that memory-boosting takes only two memory transactions to load eight request data (Fig. 3d, right). Furthermore, experiments on multiple numbers of pyramidal neurons with spines (Fig. 3e,f, S2) show that memory-boosting achieves a two fold speedup as compared to the naïve DHS.

To comprehensively test the performance of DHS with GPU memory-boosting, we select six typical neuron models and evaluate the run time of solving cable equations on massive numbers of each model (Fig. 4). We examined DHS with four threads (DHS-4) and sixteen threads (DHS-16) for each neuron, respectively. Compared to the GPU method in CoreNEURON^33^, DHS-4 and DHS-16 can speed up about 5 and 15 times, respectively (Fig. 4a). Moreover, compared to the conventional serial Hines method in NEURON running with a single-thread of CPU, DHS speeds up the simulation by 2-3 orders of magnitude (Fig. S3), while retaining the identical numerical accuracy.

**Figure 4.**
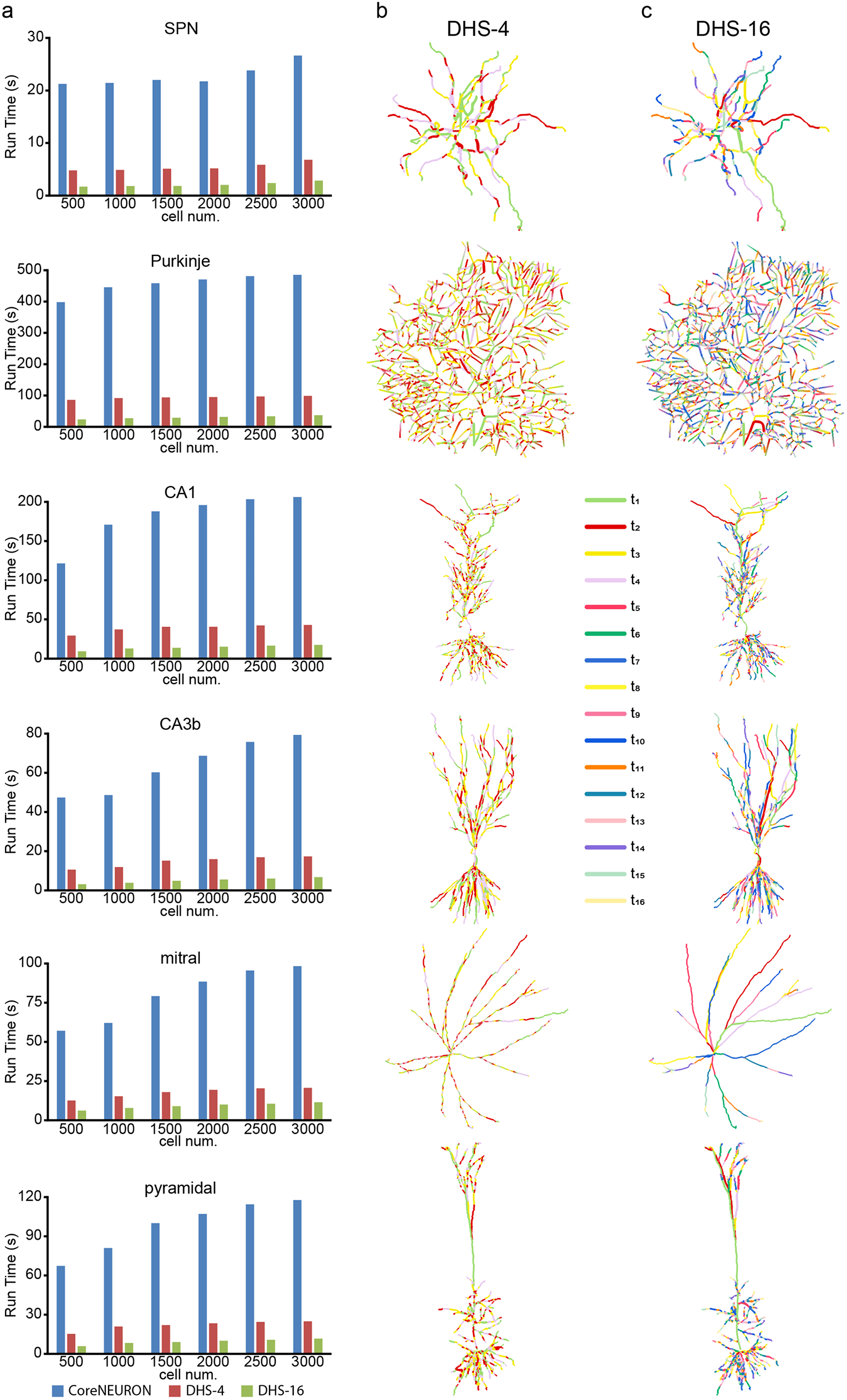
DHS enables cell-type-specific finest partition. **a**, Run time of solving equations for a 1s-simulation on GPU (dt = 0.025ms, 40000 iterations in total). CoreNEURON: parallel method used in CoreNEURON; DHS-4: DHS with four threads each neuron; DHS-16: DHS with 16 threads each neuron **b, c**, Visualization of the partition by DHS-4 and DHS-16, each color indicates a single thread. During computation, each thread switches among different branches.

### DHS Creates Cell-type-specific Optimal Partitioning

To gain insights into the working mechanism of the DHS method, we visualized the partitioning process by mapping compartments to each thread (every color presents a single thread in Fig. 4b,c). The visualization shows that a single thread frequently switches among different branches (Fig. 4b,c). Interestingly, DHS generates aligned partitions in morphologically symmetric neurons such as the striatal projection neuron (SPN) and the Mitral cell (Fig. 4b,c). By contrast, it generates fragmented partitions of morphologically asymmetric neurons like the pyramidal neurons and Purkinje cell (Fig. 4b,c), indicating that DHS splits the neural tree at individual compartment scale (i.e. tree node) rather than branch scale. This cell-type-specific fine-grained partition enables DHS to fully exploit all available threads.

In summary, DHS and memory boosting generate a theoretically proven optimal solution for solving linear equations in parallel with unprecedented efficiency. Using this principle, we built the open access DeepDendrite platform, which can be utilized by neuroscientists to implement models without any specific GPU programming knowledge.

Below, we demonstrate how we can utilize DHS and DeepDendrite in neuroscience- and AI-related tasks.

### DHS Enables Spine Level Modelling

As dendritic spines receive most of the excitatory input to cortical and hippocampal pyramidal neurons, striatal projection neurons, etc, their morphologies and plasticity are crucial for regulating neuronal excitability^10, 47-50^. However, spines are too small (∼1 μm length) to be directly measured experimentally with regard to voltage dependent processes. Thus, theoretical work is critical for the full understanding of the spine computations.

We can model a single spine with two compartments: the spine-head where synapses are located and the spine-neck that links the spine-head to dendrites^51^. The theory predicts that the very thin spine-neck (0.1-0.5 um in diameter), which electronically isolates the spine-head from its parent dendrite, thus compartmentalizing the signals generated at the spine-head^52^. However, the detailed model with fully distributed spines on dendrites (‘full-spine model’) is computationally very expensive. A common compromising solution is to modify the capacitance and resistance of the membrane by a *F*_*spines*_ factor^53^, instead of modeling all spines explicitly. Here, the *F*_*spines*_ factor aims at approximating the spine effect on the biophysical properties of the cell membrane^53^.

Inspired by the previous work of ref.^50^, we investigated how different spatial patterns of excitatory inputs formed on dendritic spines shape neuronal activities in a human pyramidal neuron model with explicitly modeled spines (Fig. 5a). Noticeably, Eyal et al. employed the *F*_*spines*_ factor to incorporate spines into dendrites while only a few activated spines were explicitly attached to dendrites (‘few-spine model’ in Fig. 5a). The value of *F*_*spines*_ in their model was computed from the dendritic area and spine area in the reconstructed data. Accordingly, we calculated the spine density from their reconstructed data to make our full-spine model more consistent with Eyal’s few-spine model. With the spine density set to 1.3/μm, the pyramidal neuron model contained about 25,000 spines without altering the model’s original morphological and biophysical properties. Further, we repeated the previous experiment protocols with both full-spine and few-spine models. We use the same synaptic input as in Eyal’s work but attach extra background noise to each sample. By comparing the somatic traces (Fig. 5b,c) and spike probability (Fig. 5d) in full-spine and few-spine models, we found that the full-spine model is much leakier than the few-spine model. In addition, the spike probability triggered by the activation of clustered spines appeared to be more nonlinear in the full-spine model (the solid blue line in Fig. 5d) than in the few-spine model (the dashed blue line in Fig. 5d). These results indicate that the conventional F-factor method may underestimate the impact of dense spine on the computations of dendritic excitability and nonlinearity.

**Figure 5.**
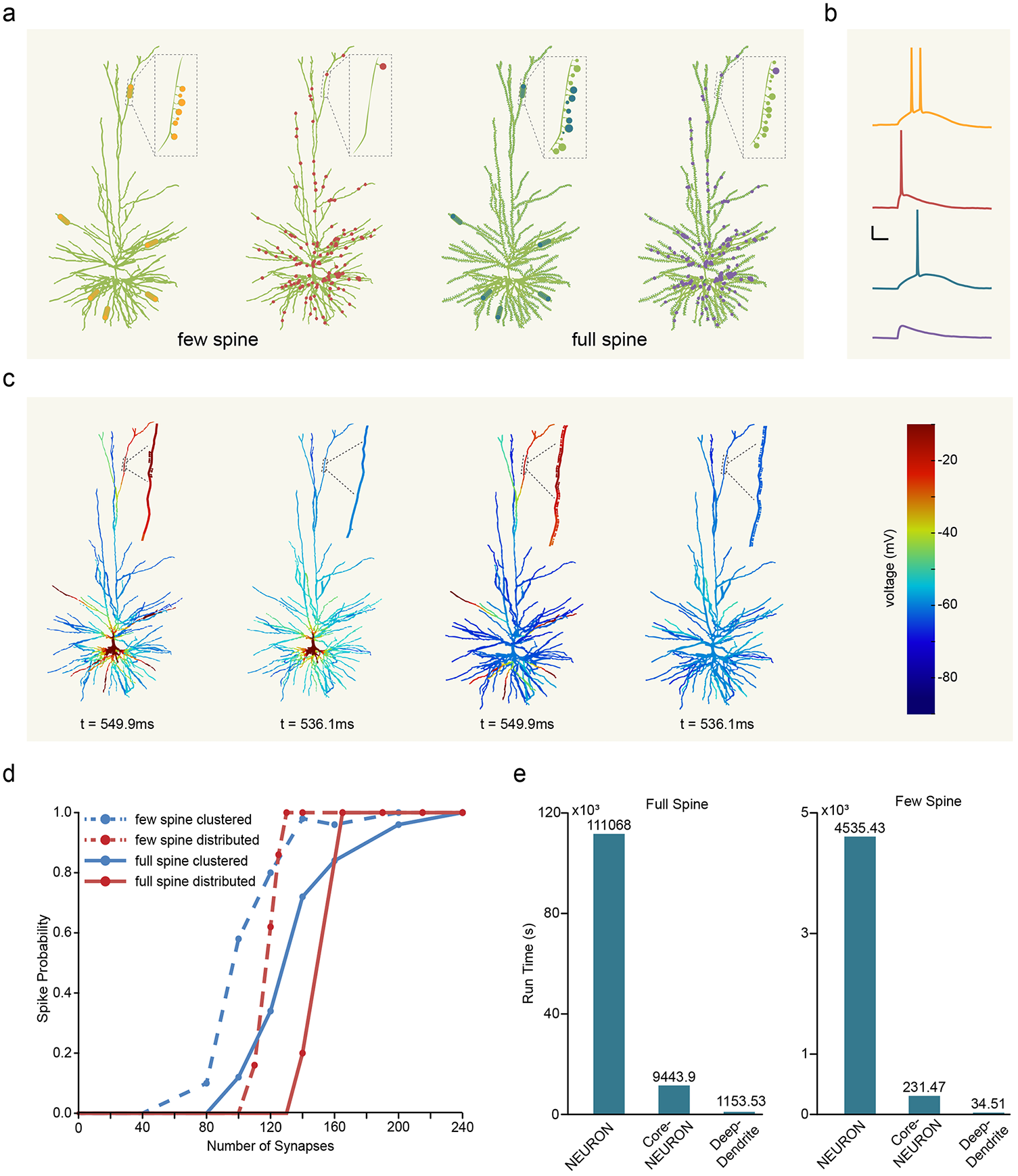
DHS enables spine level modelling. **a**, Experiment setup. We examine two major types of models: few spine model and full spine model. Few spine models (two in left) are the models that incorporated spine area globally into dendrites and only attach individual spines together with activated synapses. In full spine models (two in right), all spines are explicitly attached over whole dendrites. We explore the effects of clustered and randomly distributed synaptic inputs on the few spine model and the full spine model, respectively. **b**, Somatic voltages recorded for cases in **a**.Colors of the voltage curves e correspond to **a**, scale bar: 20ms, 20mV. **c**, Color-coded voltages during the simulation in **b** at specific time. Colors indicate the magnitude of voltage. **d**, Somatic spike probability as a function of the number of simultaneously activated synapses (as in Eyal et al.’s work) for four cases in **a**. Background noise are attached. **e**, Run time of experiments in **d** with different simulation methods. NEURON: conventional NEURON simulator run on single CPU core. CoreNEURON: CoreNEURON simulator on GPU. DeepDendrite: DeepDendrite on GPU.

In the DeepDendrite platform, both full-spine and few-spine models achieved 8 times speedup compared to CoreNEURON in the GPU platform and 100 times speedup compared to serial NEURON in the CPU platform (Fig. 5e, Table S1). Therefore, DHS method enables explorations of the dendritic excitability under more realistic anatomic conditions.

### DHS-based Learning on Detailed Dendritic Neural Networks

Deep artificial neural networks (DNNs), also called “deep learning”, have made remarkable breakthroughs in AI and exceed human performance in games and image classification tasks^2^. Because biological neural networks initially inspired DNNs, some fascinating questions remain to be answered: how does our brain realize efficient learning? Is there any learning algorithm similar to the “backprop” in DNNs? Although these questions remain open, recent work suggests that dendrites are crucial for the credit assignment in our brain, because of their enriched biophysical properties and anatomically segregated axonal projections to the dendrite from feedforward and feedback pathways. However, the earlier foundational work primarily focused on either coarse-grained dendritic network models or a single detailed compartment model. There is a lack of studies at the neural network scale with biophysically detailed models. The principal technological barriers are: 1) the computational cost of biophysically detailed networks is prohibitive; 2) the traditional simulators (NEURON, GENESIS, etc.) are not compatible with performing deep learning tasks (for details, please see Methods).

First, DHS solves the problem of the computational cost. The I/O and the learning module enable DeepDendrite to cope with the technical obstacles in learning. Thus, DeepDendrite can fully exploit detailed neural networks to explore learning algorithms and perform learning tasks (Fig. S4).

Next, we demonstrate how to implement a detailed dendritic network for a typical image classification task and utilize DHS to speed up its training procedure. We want to emphasize that here our purpose is not to build a state-of-the-art model but to explore the feasibility of training biophysically detailed networks for specific tasks. Inspired by learning rules on simplified dendritic models with only several passive compartments^17, 35^, we set up a 3-layer hybrid neural network (784×64×10), HPC-net (Human Pyramidal Cell based network). Precisely, the middle layer of HPC-net consists of 64 morphologically detailed passive human pyramidal neuron models^50^, taken from the ModelDB, access No.238347. In the HPC-net, we follow the segregation of feedforward and feedback pathways^17, 35^, while error signals are propagated from soma to dendrites for each detailed neuron to compute gradients for all the dendritic synapses (Fig. 6a, details in Method). Critically, DeepDendrite supports training with mini-batch, allowing to train dozens of copies of the network and thousands of pyramidal neuron models simultaneously in a single GPU, making the training phase converge significantly faster (Fig. 6b). With DHS, training with mini-batch becomes highly efficient, achieving a 30-fold speedup on a single GPU (NVIDIA Tesla A100) compared to the same training procedure in a 40-process-parallel NEURON on a single CPU (Intel Xeon E5-2698 v4) (Fig. 6f).

**Figure 6.**
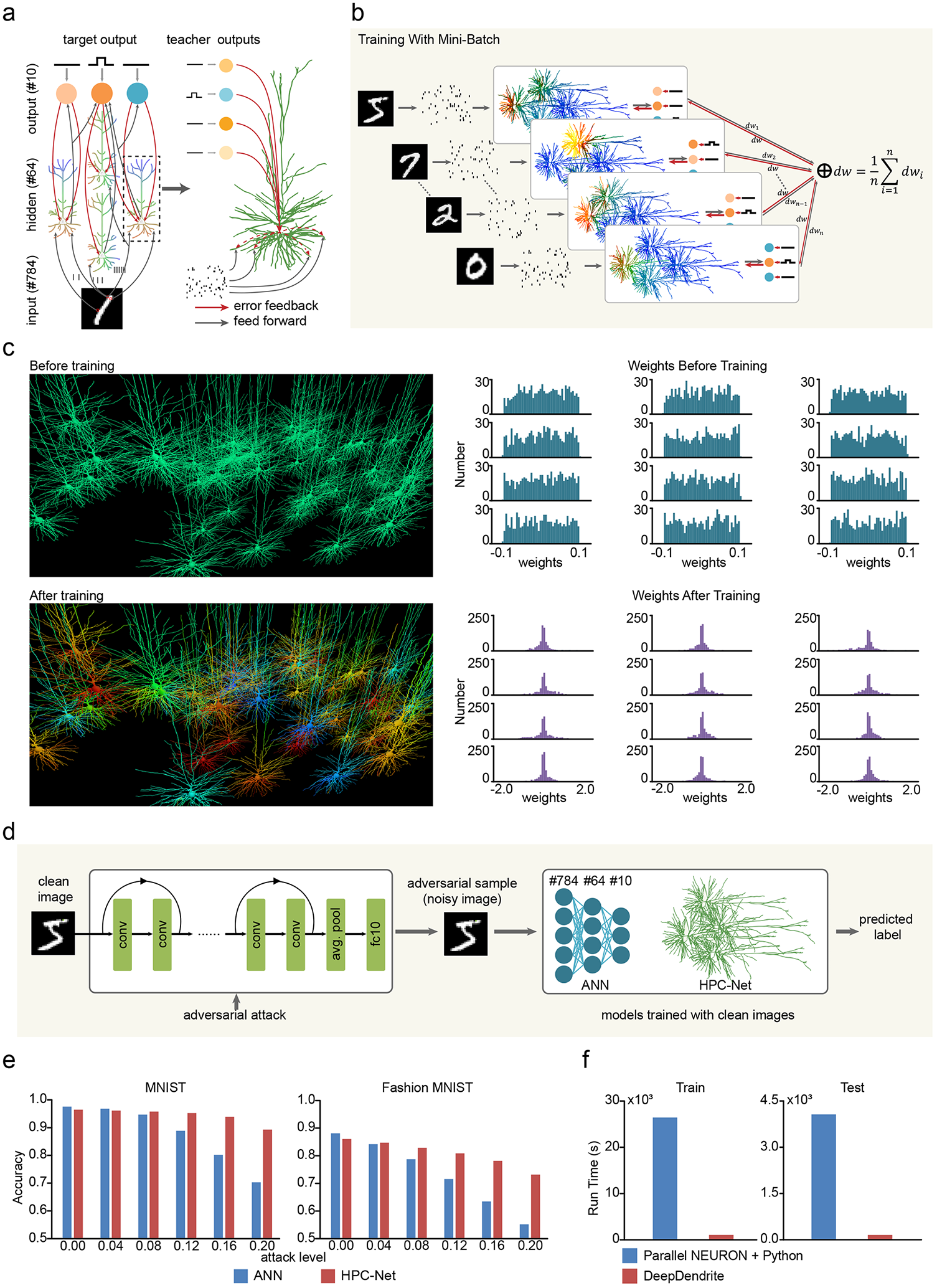
DeepDendrite enables learning with detailed neural networks. **a**, The illustration of a Human Pyramidal Cell network (HPC-Net) for image classification. Images are transformed to spike trains and fed into the network model. Learning are triggered by error signals propagated from soma to dendrites. **b**, Training with mini-batch. Multiple networks are simulated simultaneously with different images as inputs. The weights update amounts ΔW are computed as the average of ΔW_i_ from each network. **c**, Comparison of HPC-Net before and after training. Left, the visualization of hidden neuron responses to a specific input before (top) and after (bottom) training. Right, hidden layer weights (from input to hidden layer) distribution before (top) and after (bottom) training. **d**, Workflow of the defending adversarial attack experiment. We first generate adversarial samples of the test set on a 20-layer ResNet. Then use these adversarial samples (noisy images) to test the classification accuracy of models trained with clean images. **e**, Prediction accuracy of each model after training 30 epochs on adversarial samples of MNIST (left) and fashion MNIST (right) datasets. **f**, Run time of training and testing of HPC-Net, we set batch size to 16. Left, run time of training one epoch. Right, run time of testing. Parallel NEURON + Python: training and testing on multiple CPU cores, use 40-processes parallel NEURON to simulate HPC-Net, and implement I/O module and learning module with extra Python code. DeepDendrite: use DeepDendrite to train or test HPC-Net, the whole procedure of training and testing are on GPU.

To test the performance of our 3-layer HPC-Net, we next train it on typical datasets MNIST^54^ and fashion-MNIST^55^. Before training, hidden neurons are insensitive to input stimuli (Fig. 6c, upper left) as all synaptic weights are randomly distributed in -0.1 to 0.1 (Fig. 6c, upper right). After training, the distribution of synaptic weights was significantly changed (Fig. 6d, bottom right), and the detailed neurons in the hidden layer presented strong and diverse responses to input stimuli (Fig. 6d, bottom left). Furthermore, HPC-Net can achieve nearly identical prediction accuracy to a corresponding ANN (HPC-Net, 96.46% in MINIST and 86.04% in f-MNIST; ANN, 97.58% in MINIST and 88.13% in f-MNIST, Fig. 6f). For fair comparisons, the hyper-parameters of HPC-Net and the ANN, e.g. number of hidden neurons, activation function, training batch size, etc. are set the same. These data demonstrate that DeepDendrite can effectively train detailed neural networks on the image classification task.

Finally, we ask what could be the strength of detailed neural networks on image classification tasks besides that it is more biologically plausible? In deep learning, it is well known that DNNs are fragile to the intentionally designed perturbation, i.e. adversarial attacks, which the human observers can hardly perceive^56^. One intriguing hypothesis is that the dendrites and synapses protect our visual system from adversarial attacks^57^. Here we test this hypothesis with the detailed HPC-Net.

The adversarial sample is generated from intrinsic gradients of a trained network^56^. Considering the HPC-net and the ANN are two distinct networks, we adopt a popular attack technique named transferred attack to ensure that the attack is identical for both networks. We generated the adversarial samples from a third-party 20-layer ResNet^58^ rather than directly on ANN. These test adversarial samples are fed to the HPC-Net and the ANN, which are both trained with original clean images (Fig. 6d). Our results show that with the increase of attack strength, the prediction accuracy of the ANN dramatically drops to only 70.30% on MNIST and 56.47% on Fashion MNIST. In stark contrast, HPC-Net has a strong capability in defending adversarial attacks, achieving prediction accuracy of 89.30% and 73.19% under large attack strength, which is 19% and 16.72% higher than that of the ANN on MNIST and Fashion MNIST respectively (Fig. 6e). Our results are the first to show that dendrites might be crucial for biological vision robustness.

In summary, the DHS-based DeepDendrite framework enables performing learning tasks with detailed neural network models. Further, the intrinsic biophysical properties of dendrites strengthen the robustness of deep learning neural networks. Although here we only present a purely passive morphologically detailed network, such biologically detailed neural networks can be extended to train fully active detailed models with DHS. Therefore, the deep learning network will benefit from enriched biophysical properties of detailed neuron and neural network models.

## Discussion

In this work, we propose the DHS method to parallelize the computation of Hines method^38^ and we mathematically demonstrate that the DHS provides an optimal solution without any loss of precision. Next, we implement DHS on the GPU hardware platform and use GPU memory-boosting techniques to refine the DHS (Fig. 3). When simulating a large number of neurons with complex morphologies, DHS with memory-boosting achieves a 15-folds speedup (Table S1), as compared to the GPU method used in CoreNEURON^33^ and up to 1,500-folds speedup compared to serial Hines method in the CPU platform (Fig. 4, Table S1). Furthermore, we develop the DeepDendrite framework on the GPU by integrating the DHS-embedded CoreNEURON and two additional modules for learning. Thus, the DeepDendrite supports general simulations for neuroscience and for performing AI-related tasks. Finally, as demonstrations of the capacity of DeepDendrite, we present two representative applications: the first one explores the spine computations in a detailed pyramidal neuron model with 25,000 spines, while the second trains a morphologically detailed neural network for image classification tasks and tests the hypothesis of dendrites against adversarial attacks^34^. DeepDendrite can implement these tasks with unprecedented speed and therefore, significantly promotes detailed neuroscience simulations and AI explorations.

### Comparisons with previous methods

Decades of efforts have been invested in speeding up the Hines method with parallel methods. Early work mainly focuses on network-level parallelization. In network simulations, each cell independently solves its corresponding linear equations with the Hines method. Network-level parallel methods distribute a network on multiple threads and parallelize the computation of each cell groups with each thread^59, 60^. With network-level methods, we can simulate detailed networks on clusters or supercomputers^61^. In recent years, GPU has been used for detailed network simulation. Because the GPU contains massive computing units, one thread is usually assigned one cell rather than a cell group^33, 62, 63^. With further optimization, GPU-based methods achieve much higher efficiency in network simulation. However, the computation inside the cells is still serial in network-level methods, so they still cannot deal with the problem when the ‘Hines matrix’ of each cell scales large.

Cell-level parallel methods further parallelize the computation inside each cell. The main idea of cell-level parallel methods is to split each cell into several sub-blocks and parallelize the computation of those sub-blocks^25, 26^. However, typical cell-level methods (e.g. the “multi-split” method^26^) pay fewer attention to the parallelization strategy. The lack of fine parallelization strategy results in an unsatisfactory performance. To achieve higher efficiency, some studies try to obtain finer-grained parallelization by introducing extra computation operations^27, 37, 64^ or making approximation on some crucial compartments, while solving linear equations^65, 66^. These finer-grained parallelization strategies can get higher efficiency but lack sufficient numerical accuracy as in the original Hines method.

Unlike previous methods, DHS adopts the finest-grained parallelization strategy, i.e. compartment level parallelization. By modeling the problem of “how to parallelize” as a combinatorial optimization problem, DHS provides an optimal compartment-level parallelization strategy. Moreover, DHS does not introduce any extra operation or value approximation, so it achieves the lowest computational cost and retains sufficient numerical accuracy as in the original Hines’ method at the same time.

### Promoting explorations on dendritic spines and structural plasticity

Dendritic spines are the most abundant microstructures in the brain for projection neurons in cortex, hippocampus, cerebellum and the basal ganglia. As spines receive most of the excitatory inputs in the central nervous system, electrical signals generated by spines are the main driving force for large-scale neuronal activities in the forebrain and cerebellum^10, 11^. The structure of the spine, with an enlarged spine head and a very thin spine-neck—leads to surprisingly high input impedance at the spine-head, which could be up to 500 MΩ, combining experimental data and the detailed compartment modeling approach^47, 67^. Due to such high input impedance, a single synaptic input can evoke a “gigantic” EPSP (∼20 mV) at the spine-head level^47, 68^, thereby boosting NMDA currents and ion channel currents in the spine^11^. However, in the classic single detailed compartment models, all spines are replaced by the *F* coefficient modifying the dendritic cable geometries^53^. This approach may compensate for the leak currents and capacitance currents for spines. Still, it cannot reproduce the high input impedance at the spine-head, which may weaken excitatory synaptic inputs, particularly NMDA currents, thereby reducing the nonlinearity in the neuron’s input-output curve. Our modeling results are in line with this interpretation.

On the other hand, the spine’s electrical compartmentalization is always accompanied by the biochemical compartmentalization^8, 51, 69^, resulting in a drastic increase internal [Ca]^2+^,within the spine and a cascade of molecular processes involving synaptic plasticity of importance for learning and memory. Intriguingly, the biochemical process triggered by learning, in turn, remodels the spine’s morphology, enlarging (or shrinking) the spine-head, or elongating (or shortening) the spine-neck, which significantly alters the spine’s electrical capacity^69-72^. Such experience-dependent changes in spine morphology, also referred to as “structural plasticity”, have been widely observed in the visual cortex^73, 74^, somatosensory cortex^75, 76^, motor cortex ^77^, hippocampus^9^ and the basal ganglia^78^ *in vivo*. They play a critical role in motor and spatial learning, and memory formation. However, due to the computational costs, nearly all detailed network models exploit the “*F-factor*” approach to replace actual spines, and are thus unable to explore the spine functions at the system level. By taking advantage of our framework and the GPU platform, we can run a few thousands of detailed neurons models, each with tens of thousands of spines in a single GPU, while maintaining ∼100 times faster than the traditional serial method in a single CPU (Fig. 5e). Therefore, it enables us to explore of structural plasticity in large-scale circuit models across diverse brain regions.

### Promoting explorations of dendritic functions at the system level

Another critical issue is how to link dendrites to brain functions at the systems/network level. It has been well established that dendrites can perform comprehensive computations on synaptic inputs due to enriched ion channels and local biophysical membrane properties^5-7^. For example, cortical pyramidal neurons can carry out sublinear synaptic integration at the proximal dendrite but progressively shift to supralinear integration at the distal dendrite^79^. Moreover, distal dendrites can produce regenerative events such as dendritic sodium spikes, calcium spikes, and NMDA spikes/plateau potentials^6, 80^. Such dendritic events are widely observed in mice^6^ or even human cortical neurons^81^ *in vitro*, which may offer various logical operations^6, 81^ or gating functions^82, 83^. Recently, *in vivo* recordings in awake or behaving mice provide strong evidence that dendritic spikes/plateau potentials are crucial for orientation selectivity in the visual cortex^84^, sensory-motor integration in the whisker system^85, 86^, and spatial navigation in the hippocampal CA1 region^87^.

To establish the causal link between dendrites and animal (including human) patterns of behavior, large-scale biophysically detailed neural circuit models are a powerful computational tool to realize this mission. However, running a large-scale detailed circuit model of 10,000 – 100,000 neurons generally requires the computing power of supercomputers. It is even more challenging to optimize such models for *in vivo* data, as it needs iterative simulations of the models. The DeepDendrite framework can directly support many state-of-the-art large-scale circuit models^88-90^, which were initially developed based on NEURON. Moreover, using our framework, a single GPU card such as Tesla A100 could easily support the operation of detailed circuit models of up to 10,000 neurons, thereby providing carbon-efficient and affordable plans for ordinary labs to develop and optimize their own large-scale detailed models.

### Promoting dendritic learning systems with biophysically detailed models

Recent works on unraveling the dendritic roles in task-specific learning have achieved remarkable results in two directions, i.e., solving challenging tasks such as image classification dataset ImageNet with simplified dendritic networks^18^, and exploring full learning potentials on more realistic neuron^19, 20^. However, as there lies a trade-off between model size and biological detail, the increase in network scale is often sacrificed for neuron-level complexity^17, 18, 91^. Likewise, more detailed neuron models are less mathematically tractable and computationally expensive^19^.

With the help of DeepDendrite, we are able to establish our HPC-Net along with the routines for training and evaluation and reduce the total run time to an affordable amount. Although we focus only on the feasibility of training biophysically detailed networks for specific tasks, more details and insights can be brought into our HPC-net. First, as we only consider passive dendritic properties here, introducing active dendritic properties is of great importance. The work of ref.^19^ provides a fundamental framework for that and can naturally fit into our HPC-Net, since they are both based on multi-compartment modeling and computation of Hines’ method. Second, recent studies have shown that inhibitory interneurons can play an essential role in biologically plausible credit assignment. For example, local interneuron circuitry can be used along with dendritic segregation to achieve learning without assuming forward and backward phases^17, 91^. Also, disynaptic inhibition combined with short-term plasticity can decode multiplexed information streams, enabling parallel bottom-up and top-down signals^16, 18^. These features can be incorporated into our future development of deep dendritic neural networks. Overall, DeepDendrite provides first-hand feasibility for investigating the learning capability of detailed dendritic networks on the challenging tasks that require deep network architectures. It will benefit future research that seeks to bridge the field of neuroscience and modern artificial intelligence to solve credit assignments in the brain.

## Methods

### Code availability

The source code of DeepDendrite and the models in this work are available at the following link: https://disk.pku.edu.cn:443/link/AAC281DE41777C2BBB622DB2E7EF8051, or https://github.com/pkuzyc/DeepDendrite

### Simulation with DHS

CoreNEURON simulator (https://github.com/BlueBrain/CoreNeuron) uses the NEURON architecture and is optimized for both memory usage and computational speed. We implement our Dendritic Hierarchical Scheduling (DHS) method in CoreNEURON environment by modifying its source code. All models that can be simulated on GPU with CoreNEURON can also be simulated with DHS by executing the following command:

./coreneuron_exec -d */path/to/models* -e *time* –cell-permute *3* –cell-nthread *16* -- gpu

The usage options are:

|  |  |
| --- | --- |
| -d | Path containing the model data |
| -e | Simulation time (ms) |
| --cell-permute | Strategy for optimizing simulation: 1 and 2 for original strategies in CoreNEURON, **3 for DHS method** |
| --cell-nthread | Number of threads used for each cell |
| --gpu | Simulate on GPU |

### Problem formulation

#### Accuracy of the Simulation Using Cellular Level Parallel Computation

To ensure the accuracy of the simulation, we first need to define the correctness of a cellular level parallel algorithm to judge whether it will generate identical solutions compared with the proven correct serial methods, like the Hines method used in the NEURON simulation platform. Based on the theories in parallel computing^32^, a parallel algorithm will yield an identical result as its corresponding serial algorithm, *if* and *only if* the data process order in the parallel algorithm is consistent with data dependency in the serial method. The Hines method has two symmetrical phases: triangularization and back-substitution. By analyzing the serial computing Hines method^38^, we find that its data dependency can be formulated as a tree structure, where the nodes on the tree represent the compartments of the detailed neuron model. In the triangularization process, the value of each node depends on its children nodes. In contrast, during the back-substitution process, the value of each node is dependent on its parent node (Fig. 1d). Thus, we can compute nodes on different branches in parallel as their values are not dependent.

Based on the data dependency of the serial computing Hines method, we propose three conditions to make sure a parallel method will yield identical solutions as the serial computing Hines method: 1) The tree morphology and initial values of all nodes are identical to those in the serial computing Hines method; 2) In the triangularization phase, a node can be processed if and only if all its children nodes are already processed; 3) In the back-substitution phase, a node can be processed only if its parent node is already processed. Once a parallel computing method satisfies these three conditions, it will produce identical solutions as the serial computing method.

#### Computational Cost of Cellular level Parallel Computing Method

To theoretically evaluate the run time, i.e. efficiency, of the serial and parallel computing methods, we introduce and formulate the concept of computational cost as follows: given a tree *T* and *k* threads (basic computational units) to perform triangularization, parallel triangularization equals to divide the node set *V* of *T* into *n* subsets, i.e. *V* = {*V*_*1*_, *V*_*2*_, *… V*_*n*_} where the size of each subset |*V*_*i*_|≤*k*, i.e. at most *k* nodes can be processed each step since there are only *k* threads. The process of the triangularization phase follows the order: *V*_*1*_ → *V*_*2*_ → …… →*V*_*n*_, and nodes in the same subset *V*_*i*_ can be processed in parallel. So, we define |*V*| (the size of set *V*, i.e. *n* here) as the computational cost of the parallel computing method. In short, we define the computational cost of a parallel method as the number of steps it takes in the triangularization phase. Because the back-substitution is symmetrical with triangularization, the total cost of the entire solving equation phase is twice as that of the triangularization phase.

#### Mathematical Scheduling Problem

Based on the simulation accuracy and computational cost, we formulate the parallelization problem as a mathematical scheduling problem:

*Given a tree T={V, E} and a positive integer k, where V is the node-set and E is the edge set. Define partition P(V) = {V*_*1*_, *V*_*2*_, *… V*_*n*_*}*, |*V*_*i*_|≤*k, 1*≤*i*≤*n, where* |*V*_*i*_| *indicates the cardinal number of subset V*_*i*_, *i*.*e*., *the number of nodes in V*_*i*_, *and for each node v ∈V*_*i*_, *all its children nodes {c* | *c∈children(v)} must in a previous subset V*_*j*_, *where 1* ≤ *j < i. Our goal is to find an optimal partition P*^*\**^*(V) whose computational cost* |*P*^*\**^*(V)*| *is minimal*.

Here subset *V*_*i*_ consists of all nodes that will be computed at *i*^*th*^ step (Fig 2e), so |*V*_*i*_| ≤ *k* indicates that we can compute *k* nodes each step at most because the number of available threads is *k*. The restriction “*for each node v∈V*_*i*_, *all its children nodes {c* | *c ∈children(v)} must in a previous subset Vj, where 1* ≤ *j < i*” indicates that that node *v* can be processed only if all its child nodes are processed.

### DHS implementation

We aim to find an optimal way to parallelize the computation of solving linear equations for each neuron model by solving the mathematical scheduling problem above. To get the optimal partition, DHS first analyzes the topology and calculates the depth *d(v)* for all nodes *v*∈*V*. Then, the following two steps will be executed iteratively until every node *v*∈*V* is assigned to a subset: 1) find all candidate nodes and put these nodes into candidate set *Q*. A node is a candidate only if all its child nodes have been processed or it does not have any child nodes. 2) if |*Q*| ≤ *k*, i.e., the number of candidate nodes is smaller or equivalent to the number of available threads, remove all nodes in *Q* and put them into *V*_*i*_; otherwise, remove *k* deepest nodes from *Q* and add them to subset *V*_*i*_. Label these nodes as processed nodes (Fig 2d). After filling subset *V*_*i*_, go to step 1) to fill in the next subset *V*_*i+1*_.

### Proof of correctness and optimality for DHS

After applying DHS to a neural tree *T =* {*V, E*}, we get a partition *P*(*V*) = {*V*_*1*_, *V*_*2*_, … *V*_*n*_}, |*V*_*i*_| ≤ *k*, 1 ≤ *i* ≤ *n*. Nodes in the same subset *V*_*i*_ will be computed in parallel, taking *n* steps to perform triangularization and back-substitution, respectively. We then demonstrate that the reordering of the computation in DHS will result in a result identical to the serial Hines method and that its computation cost (*n)* is minimum.

#### Correctness proof

The partition *P*(*V*) obtained from DHS decides the computation order of all nodes in a neural tree. Below we demonstrate that the computation order determined by *P(V)* satisfies the correctness conditions. *P(V)* is obtained from the given neural tree *T*. Operations in DHS do not modify the tree topology and values of tree nodes (corresponding values in the linear equations), so the tree morphology and initial values of all nodes are not changed, which satisfies condition 1: the tree morphology and initial values of all nodes are identical to those in serial Hines method. In triangularization, nodes are processed from subset *V*_*1*_ to *V*_*n*_. As shown in the implementation of DHS, all nodes in subset *V*_*i*_ are selected from the candidate set *Q*, and a node can be put into *Q* only if all its child nodes have been processed. Thus the child nodes of all nodes in *V*_*i*_ are in {*V*_*1*_, *V*_*2*_, … *V*_*i-1*_}, meaning that a node is only computed after all its children have been processed, which satisfies condition 2: in triangularization, a node can be processed if and only if all its child nodes are already processed. In back-substitution, the computation order is the opposite of that in triangularization, i.e., from *V*_*n*_ to *V*_*1*_. As shown before, the child nodes of all nodes in *V*_*i*_ are in {*V*_*1*_, *V*_*2*_, … *V*_*i-1*_}, so parent nodes of nodes in *V*_*i*_ are in {*V*_*i+1*_, *V*_*i+2*_, … *V*_*n*_}, which satisfies condition 3: in back-substitution, a node can be processed only if its parent node is already processed.

#### Optimality proof

The idea of the proof is that if there is another optimal solution, it can be transformed into our DHS solution without increasing the number of steps the algorithm requires, thus indicating that the DHS solution is optimal.

For each subset *V*_*i*_ in *P(V)*, DHS moves *k* (thread number) deepest nodes from the corresponding candidate set *Q*_*i*_ to *V*_*i*_. If the number of nodes in *Q*_*i*_ is smaller than *k*, move all nodes from *Q*_*i*_ to *V*_*i*_. To simplify, we introduce *D*_*i*_, indicating the depth sum of *k* deepest nodes in *Q*_*i*_. All subsets in *P(V)* satisfy the max-depth criteria: 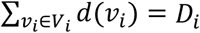. We then prove that selecting the deepest nodes in each iteration makes *P(V)* an optimal partition. If there exists an optimal partition 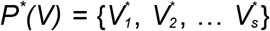 containing subsets that do not satisfy the max-depth criteria, we can modify the subsets in *P*^*\**^*(V)* so that all subsets consist of the deepest nodes from *Q* and the number of subsets (|*P*^*\**^*(V)*|) remain the same after modification.

Without any loss of generalization, we start from the first subset 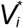 not satisfying the criteria, i.e., 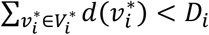. There are two possible cases that will make 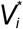 not satisfy the max-depth criteria: 1) 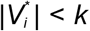 and there exists some valid nodes in *Q*_*i*_ that are not put to 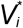; 2) 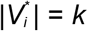 but nodes in 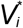 are not the *k* deepest nodes in *Q*_*i*_.

For case 1), because some candidate nodes are not put to 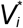, these nodes must be in the subsequent subsets. As 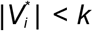, we can move the corresponding nodes from subsequent subsets to 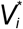, which will not increase the number of subsets and make 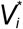 satisfy the criteria (Fig. S6b, top). For case 2), 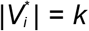, these deeper nodes that are not moved from the candidate set *Q*_*i*_ into 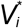 must be added to subsequent subsets (Fig. S6b, bottom). These deeper nodes can be moved from subsequent subsets to 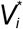 through the following method. Assume that after filling 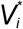, *v* is picked and one of the *k*^*th*^ deepest nodes *v’* is still in *Q*_*i*_, thus *v’* will be put into a subsequent subset 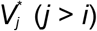. We first move *v* from 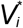 to 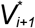, then modify subset 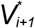 as follows: if 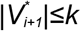 and none of the nodes in 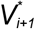 is the parent of node *v*, stop modifying the latter subsets. Otherwise, modify 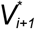 as follows: if the parent node of *v* is in 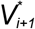, move this parent node to 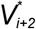; else move the node with minimum depth from 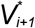 to 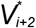. After adjusting 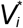, modify subsequent subsets 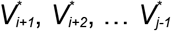 with the same strategy. Finally, move *v’* from 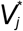 to 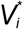.

With the modification strategy described above, we can replace all shallower nodes in 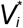 with the *k*^*th*^ deepest node in *Q*_*i*_ and keep the number of subsets, i.e. | *P*^*\**^*(V)*| the same after modification. We can modify the nodes with the same strategy for all subsets in *P*^*\**^*(V)* that do not contain the deepest nodes. Finally, all subsets 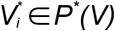 can satisfy the max-depth criteria, and |*P*^*\**^*(V)*| does not change after modifying.

In conclusion, DHS generates a partition *P(V)*, and all subsets *V*_*i*_∈*P(V)* satisfy the max-depth condition: 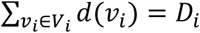. For any other optimal partition *P*^*\**^*(V)* we can modify its subsets to make its structure the same as *P(V)*, i.e., each subset consists of the deepest nodes in the candidate set, and keep |*P*^*\**^*(V)*| the same after modification. So, the partition *P(V)* obtained from DHS is one of the optimal partitions.

### GPU implementation and memory boosting

To achieve high memory throughput, GPU utilizes the memory hierarchy of 1) global memory, 2) cache, 3) register, where global memory has large capacity but low throughput, while registers have low capacity but high throughput. We aim to boost memory throughput by leveraging the memory hierarchy of GPU.

GPU employs SIMT (Single-Instruction, Multiple-Thread) architecture. Warps are the basic scheduling units on GPU (a warp is a group of 32 parallel threads). A warp executes the same instruction with different data for different threads^45^. Correctly ordering the nodes is essential for this batching of computation in warps, to make sure DHS obtains identical results as the serial Hines method. When implementing DHS on GPU, we first group all cells into multiple warps based on their morphologies. Cells with similar morphologies are grouped in the same warp. We then apply DHS on all neurons, assigning the compartments of each neuron to multiple threads. Because neurons are grouped into warps, the threads for the same neuron are in the same warp. Therefore, the intrinsic synchronization in warps keeps the computation order consistent with the data dependency of the serial Hines method. Finally, threads in each warp are aligned and rearranged according to the number of compartments.

When a warp loads pre-aligned and successively-stored data from global memory, it can make full use of the cache, which leads to high memory throughput, while accessing scatter-stored data would reduce memory throughput. After compartments assignment and threads rearrangement, we permute data in global memory to make it consistent with computing orders so that warps can load successively-stored data when executing the program. Moreover, we put those necessary temporary variables into registers rather than global memory. Registers have the highest memory throughput, so the use of registers further accelerates DHS.

### Explore neuronal excitability with full spine models

#### Biophysical model

We used the published human pyramidal neuron^50^. The membrane capacitance *c*_*m*_ = 0.44 μF/cm^2^, membrane resistance *r*_*m*_ = 48300 Ω cm^2^ and axial resistivity *r*_*a*_ = 261.97 Ω cm. In this model, all dendrites were modeled as passive cables while somas were active. The leak reversal potential *E*_*l*_ = -83.1 mV. Ion channels such as Na^+^ and K^+^ were inserted on soma and axon, and their reversal potentials were *E*_*Na*_ = 67.6 mV, *E*_*K*_ = -102 mV respectively. All these specific parameters were set the same as in the model of ref.^50^, for more details please refer to the published model (ModelDB, access No.238347).

In the few-spine model, the membrane capacitance and maximum leak conductance of the dendritic cables 60 μm away from soma were multiplied by a *F*_*spine*_ factor to approximate dendritic spines. In this model, *F*_*spine*_ was set to 1.9. Only the spines that receive synaptic inputs were explicitly attached to dendrites.

In the full-spine model, all spines were explicitly attached to dendrites. We calculated the spine density with the reconstructed neuron in ref^50^. The spine density was set to 1.3/μm, and each cell contained 24994 spines on dendrites 60 μm away from the soma.

The morphologies and biophysical mechanisms of spines were the same in few-spine and full-spine models. The length of the spine neck *L*_*neck*_ = 1.35 μm and the diameter *D*_*neck*_ = 0.25 μm, whereas the length and diameter of the spine head were 0.944 μm, i.e., the spine head area was set to 2.8 μm^2^. Both spine neck and spine head were modeled as passive cables, with the reversal potential *E*_*l*_ = -86 mV. The specific membrane capacitance, membrane resistance, and axial resistivity were the same as those for dendrites.

#### Synaptic inputs

We investigated neuronal excitability for both distributed and clustered synaptic inputs. All activated synapses were attached to the terminal of the spine head. For distributed inputs, all activated synapses were randomly distributed on all dendrites. For clustered inputs, each cluster consisted of 20 activated synapses that were uniformly distributed on a single randomly-selected compartment. All synapses were activated simultaneously during the simulation.

AMPA-based and NMDA-based synaptic currents were simulated as in Eyal et al.’s work. AMPA conductance was modeled as a double-exponential function and NMDA conduction as a voltage-dependent double-exponential function. For AMPA model, the specific *τ*_*rise*_ and *τ*_*decay*_ were set to 0.3 and 1.8 ms. For NMDA model, *τ*_*rise*_ and *τ*_*decay*_ were set to 8.019 and 34.9884 ms, respectively. The maximum conductance of AMPA and NMDA were 0.73 nS and 1.31 nS.

#### Background Noise

We attached background noise to each cell to simulate a more realistic environment. Noise patterns were implemented as Poisson spike trains with a constant rate of 1.0 Hz. Each pattern started at *t*_*start*_ =10ms and lasted until the end of the simulation. We generated 400 noise spike trains for each cell and attached them to randomly-selected synapses. The model and specific parameters of synaptic currents were the same as those in ***Synaptic Inputs***, except that the maximum conductance of NMDA was uniformly distributed from 1.57 to 3.275, resulting in a higher AMPA to NMDA ratio.

#### Explore neuronal excitability

We investigated the spike probability when multiple numbers of synapses were activated simultaneously. For distributed inputs, we tested 14 cases, from 0 to 240 activated synapses. For clustered inputs, we tested 9 cases in total, activating from 0 to 12 clusters respectively. Each cluster consisted of 20 synapses. For each case in both distributed and clustered inputs, we calculated the spike probability with 50 random samples. Spike probability was defined as the ratio of fired neuron number to the total sample number. All 1150 samples were simulated simultaneously on our DeepDendrite platform, reducing the simulation time from days to minutes.

### Perform AI task with DeepDendrite platform

Conventional simulators lack two functionalities important to modern AI tasks: 1) processing multiple stimuli samples in a batch-like manner and 2) updating synaptic weights during simulation without manual intervention. Here we present the DeepDendrite platform, which supports both biophysical simulating and performing deep learning tasks with detailed dendritic models.

DeepDendrite consists of three modules: 1) I/O module; 2) DHS-based simulating module; 3) learning module. When training a biophysically detailed model to perform learning tasks, users first define the learning rule, then feed all training samples to the detailed model for learning. In each step during training, the I/O module picks a specific stimulus and its corresponding teacher signal (if necessary) from all training samples and attaches the stimulus to the network model. Then, DHS-based simulating module initializes the model and starts the simulation. After simulation, the learning module updates all synaptic weights according to the difference between model responses and teacher signals. After training, the learned model can achieve performance comparable with ANN. The testing phase is similar to training, except that all synaptic weights are fixed.

### Image classification with the biophysically detailed network model

Image classification is a typical task in the artificial intelligence (AI) field. In this task, a model should learn to recognize the content in a given image and output the corresponding label. Here we present the HPC-Net, a network consisting of detailed human pyramidal neuron models that can learn to perform image classification tasks by utilizing the DeepDendrite platform.

#### HPC-Net model

HPC-Net has three layers, i.e., an input layer, a hidden layer, and an output layer. The neurons in the input layer receive spike trains converted from images as their input. Hidden layer neurons receive the output of input layer neurons and deliver responses to neurons in the output layer. The responses of the output layer neurons are taken as the final output of HPC-Net. Neurons between adjacent layers are fully connected.

For each image stimulus, we first convert each normalized pixel to a homogeneous spike train. The interval *int(x, y)* between two spikes is constant in each spike train and is determined by the pixel value *p(x, y)* as shown in Equation 1.

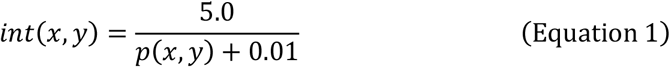

In our experiment, the simulation for each stimulus lasted 50 ms. All spike trains started at 9 ms and lasted until the end of the simulation. Then we attached all spike trains to the input layer neurons in a one-to-one manner. The synaptic current triggered by the spike arriving at time *t*_*0*_ is given by

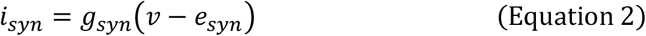

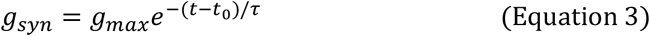

where *v* is the voltage of post synapse, the reversal potential *e*_*syn*_ = 1mV, the maximum synaptic conductance *g*_*max*_=0.05uS, and the time constant *τ* = 0.5ms.

Neurons in the input layer were modeled with a passive single-compartment model. The specific parameters were set as follows: membrane capacitance *c*_*m*_ = 1.0 μF/cm^2^, membrane resistance *r*_*m*_ = 1e4 Ω cm^2^, axial resistivity *r*_*a*_ = 100 Ω cm, reversal potential of passive compartment *E*_*l*_ = 0mV.

The hidden layer contains a group of human pyramidal neuron models, receiving the somatic voltages of input layer neurons. The morphology was from ref.^50^, and all neurons were modeled with passive cables. The specific membrane capacitance *c*_*m*_ = 1.5 μF/cm^2^, membrane resistance *r*_*m*_ = 48300 Ω cm^2^, axial resistivity *r*_*a*_ = 261.97 Ω cm, and the reversal potential of all passive cables *E*_*l*_ = -86 mV. Input neurons made multiple connections to randomly-selected locations on the dendrites of hidden neurons. The synaptic current activated by the *i*^*th*^ input neuron on dendrites is defined as in Equation 4, where *g*_*h*_ is a constant value for scaling, 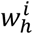 is the synaptic weight and 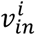 is the somatic voltage of the *i*^*th*^ input neuron.

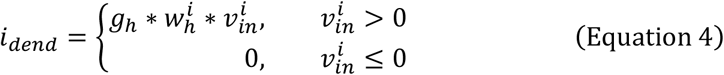

Neurons in the output layer were also modeled with a passive single-compartment model, and each neuron took the somatic voltages of hidden neurons as input. All specific parameters were set the same as those of the input neurons. Synaptic currents activated by hidden neurons are also in the form of Equation 4.

#### Image classification with HPC-Net

For each input image stimulus, we first normalized all pixel values to 0.0-1.0. Then we converted normalized pixels to spike trains and attached them to input neurons. Somatic voltages of the output neurons are used to compute the predicted probability of each class, as shown in Equation 5, where *p*_*i*_ is the probability of *i*^*th*^ class predicted by the HPC-Net, 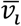 is the mean somatic voltage during 20ms to 50ms of the *i*^*th*^ output neuron, and *C* indicates the number of classes, which equals the number of output neurons. The class with maximum predicted probability is the final classification result. In this paper, we built the HPC-Net with 784 input neurons, 64 hidden neurons and 10 output neurons.

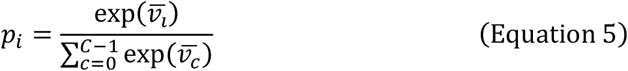

#### Learning on HPC-Net

Inspired by previous work^35^, we use a gradient-based learning rule to train our HPC-Net to perform the image classification task. The loss function we use here is cross-entropy, given in Equation 6, where *p*_*i*_ is the predicted probability for class *i, y*_*i*_ indicates the actual class the stimulus image belongs to, *y*_*i*_=1 if input image belongs to class *i*, and *y*_*i*_=0 if not.

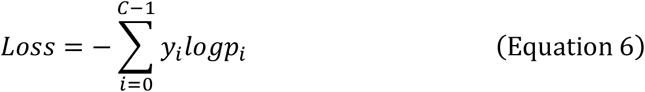

When training HPC-Net, we compute the update for weight *W*_*ijk*_ (the synaptic weight of the *k*^*th*^ synapse connecting neuron *i* to neuron *j*) at each timestep. After the simulation of each image stimulus, *W*_*ijk*_ is updated as shown in Equation 7:

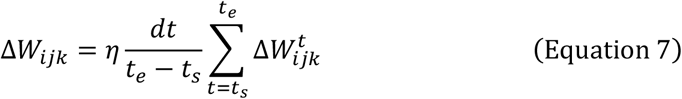

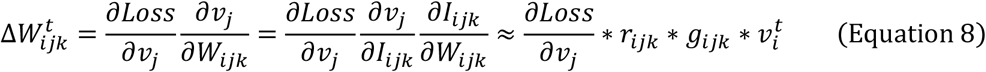

Here *η* is the learning rate, 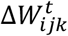 is the update value at time *t, v*_*j*_, *v*_*i*_ are somatic voltages of neuron *i* and *j* respectively, *I*_*ijk*_ is the *k*^*th*^ synaptic current activated by neuron *i* on neuron *j, g*_*ijk*_ its synaptic conductance, *r*_*ijk*_ is the transfer resistance between the *k*^*th*^ connected compartment of neuron *i* on neuron *j*’s dendrite to neuron *j*’s soma, *t*_*s*_ = 30 ms, *t*_*e*_ = 50 ms are start time and end time for learning respectively. For output neurons, the error term 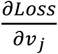 can be computed as shown in Equation 9. For hidden neurons, the error terms are calculated from the error terms in the output layer, given in Equation 10.

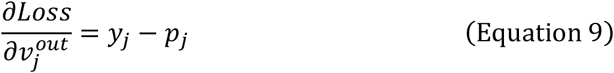

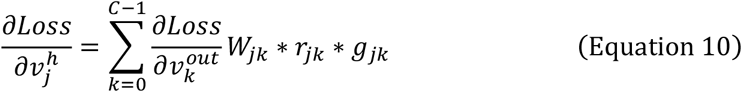

Since all output neurons are single-compartment, *r*_*ik*_ equals to the input resistance of the corresponding compartment, *r*_*kk*_.

Training with mini-batch is a typical method in deep learning for achieving higher prediction accuracy and accelerating convergence in training. DeepDendrite also supports mini-batch training. When training HPC-Net with mini-batch size *N*_*batch*_, we make *N*_*batch*_ copies of HPC-Net. During training, each copy is fed with a different training sample from the batch. DeepDendrite first computes the weight update for each copy separately. After all copies in the current training batch are done, the average weight update is calculated and weights in all copies are updated by this same amount.

#### Defense adversarial attack with HPC-Net

To demonstrate the robustness of HPC-Net, we tested its prediction accuracy on adversarial samples and compared it with an analogous ANN (one with the same 784-64-10 structure and ReLU activation). We first trained HPC-Net and ANN with the original training set (original clean images). Then we added adversarial noise to the test set and measured their prediction accuracy on the noisy test set. We used the foolbox (https://github.com/bethgelab/foolbox) to generate adversarial noise with FGSM method^56^. ANN was trained with Tensorflow (https://tensorflow.google.cn), and HPC-Net was trained with our DeepDendrite. For fairness, we generated adversarial noise on a significantly different network model, a 20-layer ResNet^58^. The noise level ranged from 0.02 to 0.2. We experimented on two typical datasets, MNIST^54^ and Fashion MNIST^55^. Results show that the prediction accuracy of HPC-Net is 19% and 16.72% higher than that of the analogous ANN, respectively.

## Acknowledgments

The authors thank Rita Zhang, Daochen Shi and members at NVIDIA for the technical help. This work is supported by National Key R&D Program of China (No. 2020AAA0130400).

