## Supplementary tables and figures for "A GPU-based computational framework that bridges Neuron simulation and Artificial Intelligence"

Table S1 Summary of speedup referred in our experiments

| Ours | Phase | Compared method | Model | SpeedUp |
|---|---|---|---|---|
| DHS | solving equations | CoreNEURON method (GPU) | pyramidal | 10.08-11.35 |
| | | | purkinje | 13.11-17.00 |
| | | | CA3b | 11.76-15.00 |
| | | | CA1 | 11.78-13.63 |
| | | | mitral | 7.85-9.14 |
| | | | SPN | 9.39-12.56 |
| | | serial Hines method (CPU) | pyramidal | 174.11-953.65 |
| | | | purkinje | 285.67-1589.42 |
| | | | CA3b | 145.43-800.48 |
| | | | CA1 | 238.41-1162.62 |
| | | | mitral | 127.85-713.91 |
| | | | SPN | 67.10-270.73 |
| DeepDendrite | simulation | serial NEURON (CPU) | full spine model | 96.37 |
| | | CoreNEURON (GPU) | full spine model | 8.19 |
| | | serial NEURON (CPU) | few spine model | 131.42 |
| | | CoreNEURON (GPU) | few spine model | 6.71 |
| | training | 40-process parallel NEURON (CPU) | HPC-Net | 25.39 |
| | testing | 40-process parallel NEURON (CPU) | HPC-Net | 26.54 |

CoreNEURON version: 0.14, NEURON version: 7.6.2; CPU: Intel Xeon E5-2698 v4, GPU: NVIDIA Tesla A100

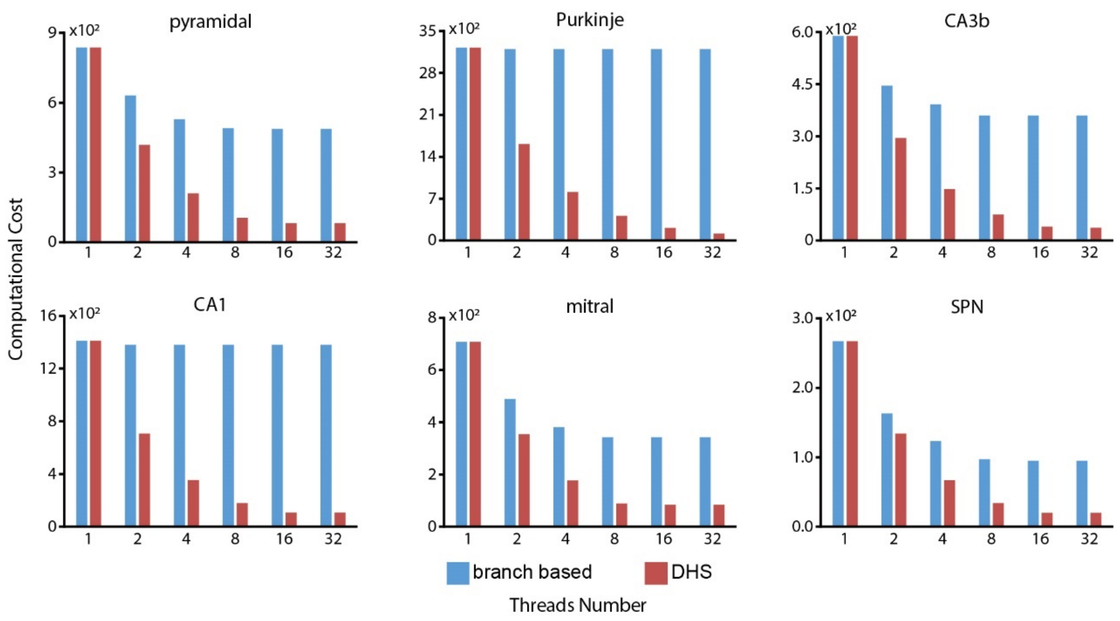


**Figure S1 DHS outperforms branch based method in computational cost.** DHS achieves low computational cost on various types of neurons while the costs of branch based method are much higher. The cost of branch based method remains high despite the growth of thread number, and in some cases (e.g. Purkinje cell and CA1 cell) only little gain on cost is achieved.

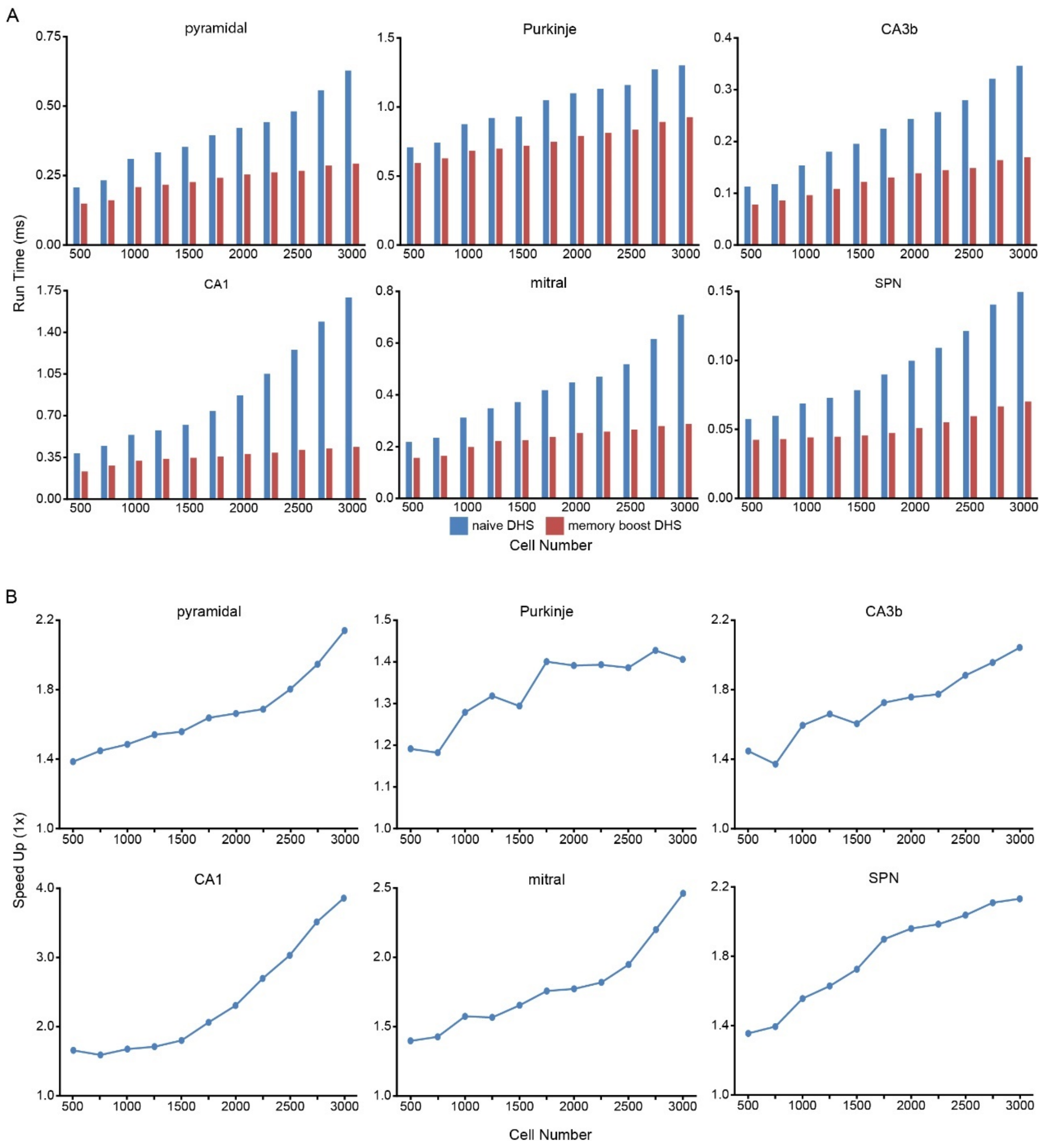


**Figure S2 GPU memory boosting further speeds up DHS.** (A) GPU memory boosting reduce run time of DHS on various types of neurons. (B) With GPU memory boosting, DHS becomes 1.2-3.8 times faster on different types of neurons

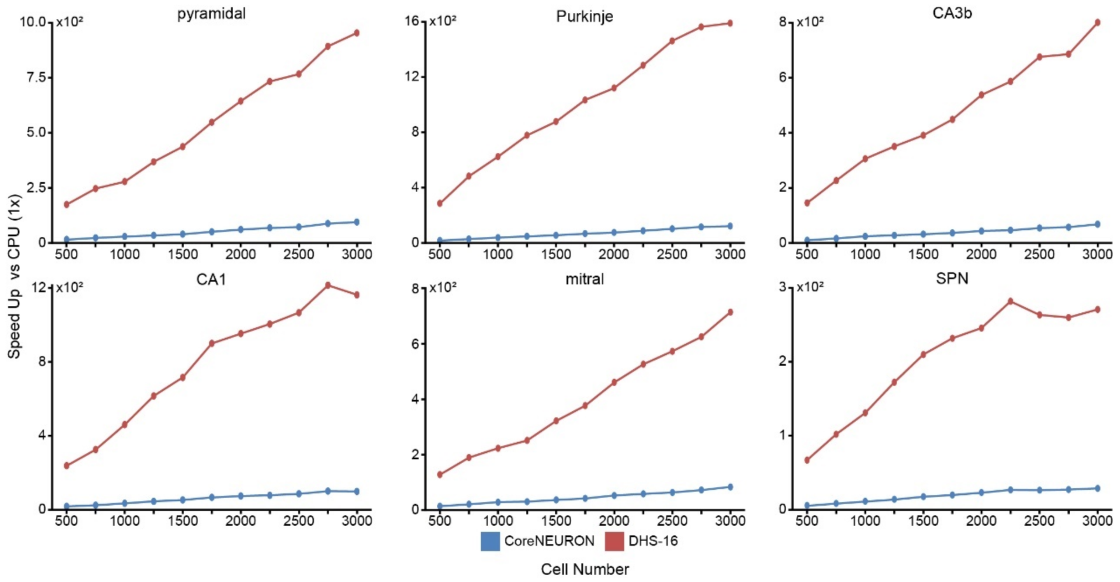


**Figure S3 DHS achieves 2-3 orders of magnitude speedup compared to the serial Hines method on CPUs.** On the test neurons used in this study, the GPU parallel method used in CoreNEURON achieves speed up of 2.5-120 folds compared to serial Hines method on CPU. DHS method is 7-17 times faster than the method in CoreNEURON, and achieves speed up of 60-1500 folds compared to serial Hines method on CPU.

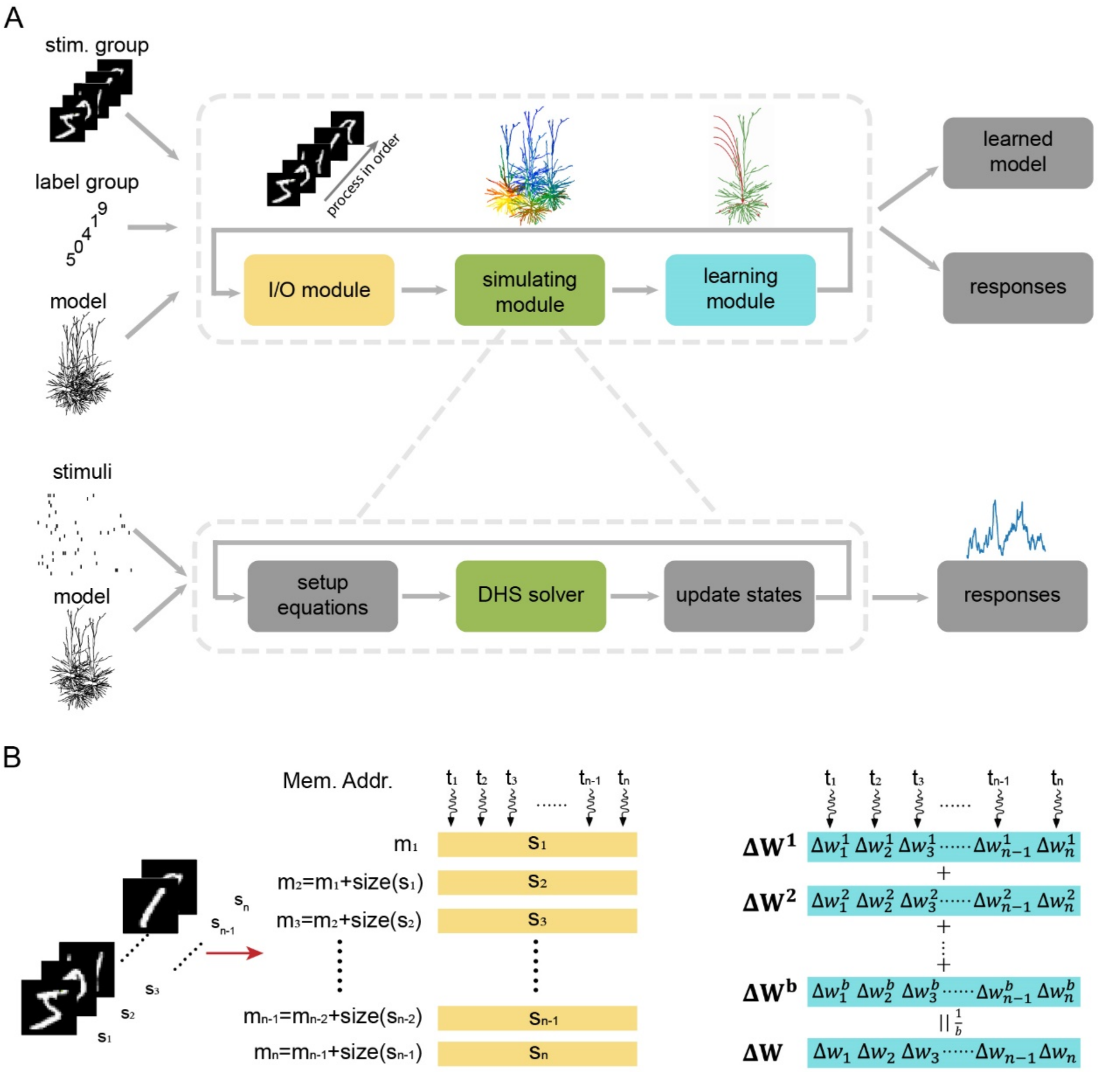


C

**Algorithm 1** Training on DeepDendrite

**Input** Model M, Training Set $TrainSet$, Training Set size $N$, batch size $b$, iteration number $N_{iter}$

**Output** Weights after training $\mathbf{W}$

**function** GPUTRAIN(M, $TrainSet, N, b, N_{iter}$)
  $istim = 0$
  $iter = 0$
  **while** $iter < N_{iter}$ **do**
    $\mathbf{s}, \mathbf{l} = TrainSet[istim : (istim + b)\%N]$
    Attach stimuli $\mathbf{s}$ and labels $\mathbf{l}$ to the model M
    Simulate to get the responses of all neurons
    Compute $\Delta \mathbf{W}_1, \dots \Delta \mathbf{W}_b$
    Set $\Delta \mathbf{W} = mean(\Delta \mathbf{W}_1, \dots \Delta \mathbf{W}_b)$
    Update all synaptic weights $\mathbf{W} = \mathbf{W} + \eta \Delta \mathbf{W}$
    $istim = (istim + b)\%N$
    $iter += 1$
  **end while**
**end function**

**Algorithm 2** Testing on DeepDendrite

**Input** Model M, Test Set $TestSet$, Set size $N$, batch size $b$

**Output** Responses of the model $\mathbf{Y}_{pred}$

**function** GPUTEST(M, $TestSet, N, b$)
  $istim = 0$
  **while** $istim < N$ **do**
    $\mathbf{s}, \mathbf{l} = TestSet[istim : istim + b]$
    Attach stimuli $\mathbf{s}$ to the model
    Simulate to get all neural responses $y_1, y_2 \dots y_b$
    Set $\mathbf{Y}_{pred}[istim : istim + b] = \{y_1, y_2, \dots, y_b\}$
    $istim = (istim + b)\%N$
  **end while**
**end function**

**Figure S4 The framework and implementation of DeepDendrite. (A) DeepDendrite framework.** DeepDendrite consists of three modules: I/O module, simulating module and learning module. During learning, DeepDendrite takes the detailed model and all training samples as input, and save the model after learning. During each iteration in training, I/O module picks specific stimuli from all training samples and attach them to the detailed model. Then simulating module starts simulation and gets the responses of the detailed model. After simulation, learning module updates synaptic weights according to the network responses and the target signal. (B) GPU implementation and optimization of I/O module (left) and learning module (right). (C) The procedure of training (left) and testing (right) when performing AI tasks with detailed network model on DeepDendrite.

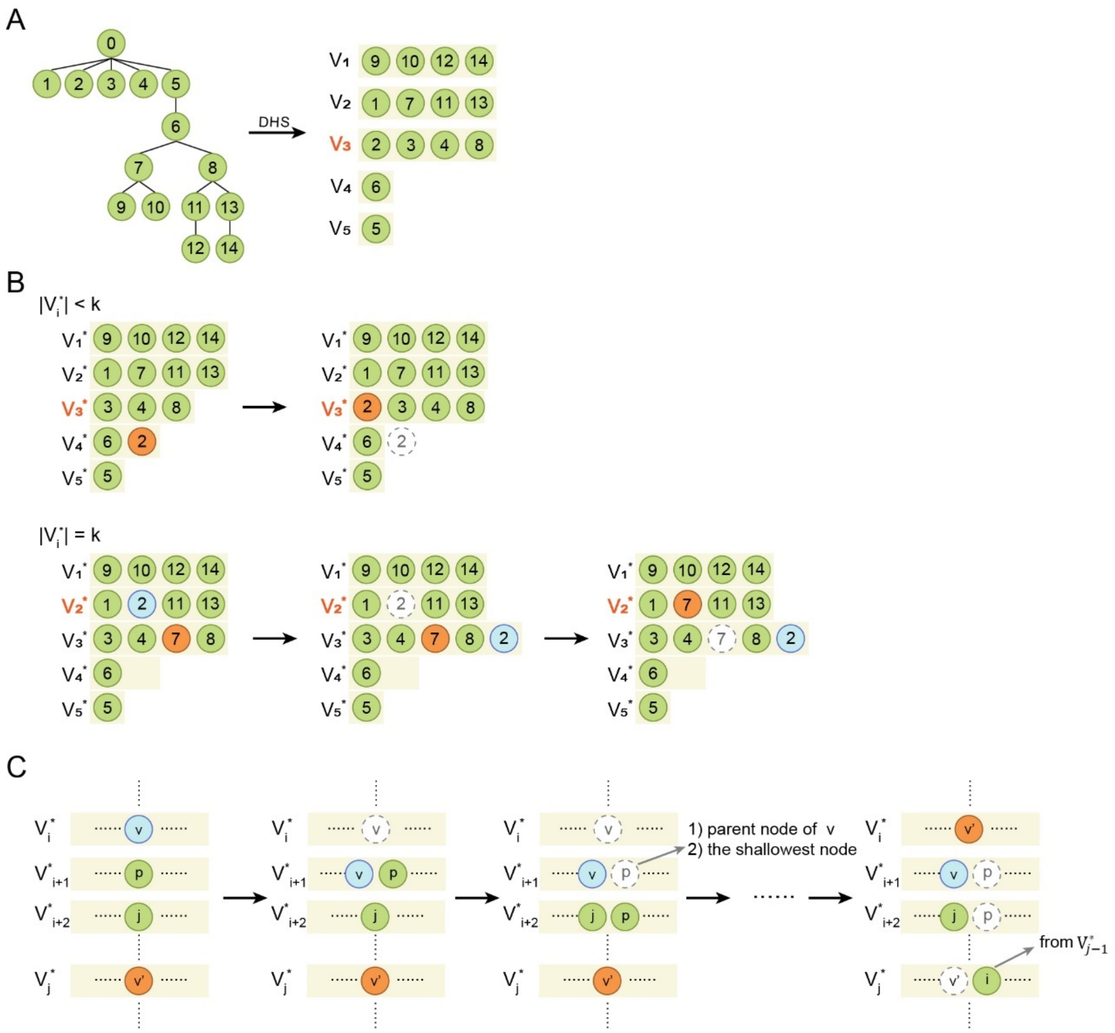


**Figure S5 Strategies for modifying subset $V_i^*$ to make it satisfy the max-depth criteria.** (A) partition generated by DHS (B) A simple example showing how to modify subset $V_i^*$. (C) The general case when $|V_i^*| = k$
